# Fetal estrogens are not involved in sex determination but critical for early ovarian differentiation

**DOI:** 10.1101/2020.12.13.419770

**Authors:** Geneviève Jolivet, Nathalie Daniel-Carlier, Erwana Harscoёt, Eloïse Airaud, Aurélie Dewaele, Cloé Pierson, Frank Giton, Laurent Boulanger, Nathalie Daniel, Béatrice Mandon-Pépin, Maёlle Pannetier, Eric Pailhoux

## Abstract

AROMATASE, encoded by the *CYP19A1* gene, is the cytochrome enzyme responsible for the synthesis of estrogens in vertebrates. In most mammals a peak of expression of the *CYP19A1* gene occurs in the fetal XX gonad when sexual differentiation starts up. To elucidate the role of this peak, we produced 3 lines of TALEN genetically edited *CYP19A1* KO rabbits, that were void of any production of estradiol. All KO XX rabbits developed as females, with aberrantly small sized ovaries at adulthood, an almost empty reserve of primordial follicles and very few large antrum follicles. Ovulation never occurred. Our histological, immunohistological and transcriptomic analyses showed that the surge of estradiol in the XX fetal rabbit gonad is dispensable for its determination as an ovary, or for meiosis. However, it is mandatory for the high proliferation and differentiation of both somatic and germ cells, and consequently for the establishment of the ovarian reserve.

---

In Vertebrates, AROMATASE is the unique enzyme responsible for the synthesis of estrogens. It converts specifically three androgenic substrates (androstenedione, testosterone and 17alpha-testosterone) to estrone, estradiol, and estriol respectively. It is the product of the *CYP19A1* gene, which is the unique member of its gene family. Although the *CYP19A1* gene is commonly known to be mainly expressed by gonads in adults, a physiologically significant expression has already been reported in numerous tissues or organs such as bones, adipose tissue, brain or placenta (the expression in the latter depending on the species, see for review in (Conley and Hinshelwood 2001) and references herein).

In mammals, the expression of the *CYP19A1* gene begins early at around implantation in the embryo in rabbits (Dickmann *et al.* 1975; George and Wilson 1978b), pigs and ruminants (Gadsby *et al.* 1980). A second burst of expression was reported later in fetal life in the ovaries and not in testes shortly after the sex determination process. This was described in rabbits (George *et al.* 1978; George and Wilson 1978b; Gondos *et al.* 1983; Daniel-Carlier *et al.* 2013), sheep (Mauleon *et al.* 1977; Payen *et al.* 1996; Torley *et al.* 2011), goats (Pannetier *et al.* 2006), bovines (Shemesh 1980; Ross *et al.* 2009; Garverick *et al.* 2010) and humans (George and Wilson 1978a) by RNA measurements or by estrogen assays in gonads or gonadal culture media. This peak is transitory and spans until meiosis initiation. Later on, *CYP19A1* gene expression increases progressively with follicle differentiation. It culminates in antrum follicles, the mural granulosa cells being the site of expression. By the past, the physiological significance of this fetal peak of expression has been searched in the rabbit species. Namely, George et al (George *et al.* 1978) have supposed that the local production of estrogens in the XX gonad was a key event for the initiation of the differentiation of the female fetal gonad but this hypothesis has still not been proven. In birds (Elbrecht and Smith 1992; Vaillant *et al.* 2001) and fishes (Guiguen *et al.* 2010), it was shown that estrogens are major determinants of the ovary orientation of the undifferentiated gonad in fetal or larval life. The question was thus to investigate the role of the fetal peak of estrogen in the mammalian gonad and more specifically to search whether in mammals it is mandatory for ovary differentiation.

To reach this aim, we decided to suppress the expression of the *CYP19A1* gene in a mammal species other than the mice. Indeed, rodents represent some of the few mammalian species without detectable peak of expression at this developmental stage in the fetal XX gonad i.e. before meiosis. Thus, thanks to the TALEN methodology already shown to be efficient for gene targeting in the rabbit species (Lee *et al.* 2018; Peyny *et al.* 2020), we produced *CYP19A1* knocked-out (KO) rabbits completely devoid of estrogens secretion. Here we report the phenotype of the XX *CYP19A1* KO (*ARO^-/-^*) female rabbits with severe ovarian developmental defects. A series of histological and genetical analyses was thus carried out from the early stages of gonadal differentiation to investigate the mechanisms involved.

## Results

### Three aromatase mutant rabbit lines which do not produce estradiol were generated

The TALEN subunits were designed to generate *InDel* mutations through non-homologous end joining near the initiation site of translation of the *CYP19A1* gene as described in Figure 1. After transferring 67 injected embryos in 3 recipient female rabbits, 6 founders (2 males and 4 females) harboring mutant alleles were characterized from 11 neonates. Each founder F0 was mated with a wild type counterpart in order to transmit a mutant allele to the descendants and generate mutant lines carrying one mutated allele per line.

**Figure 1:**
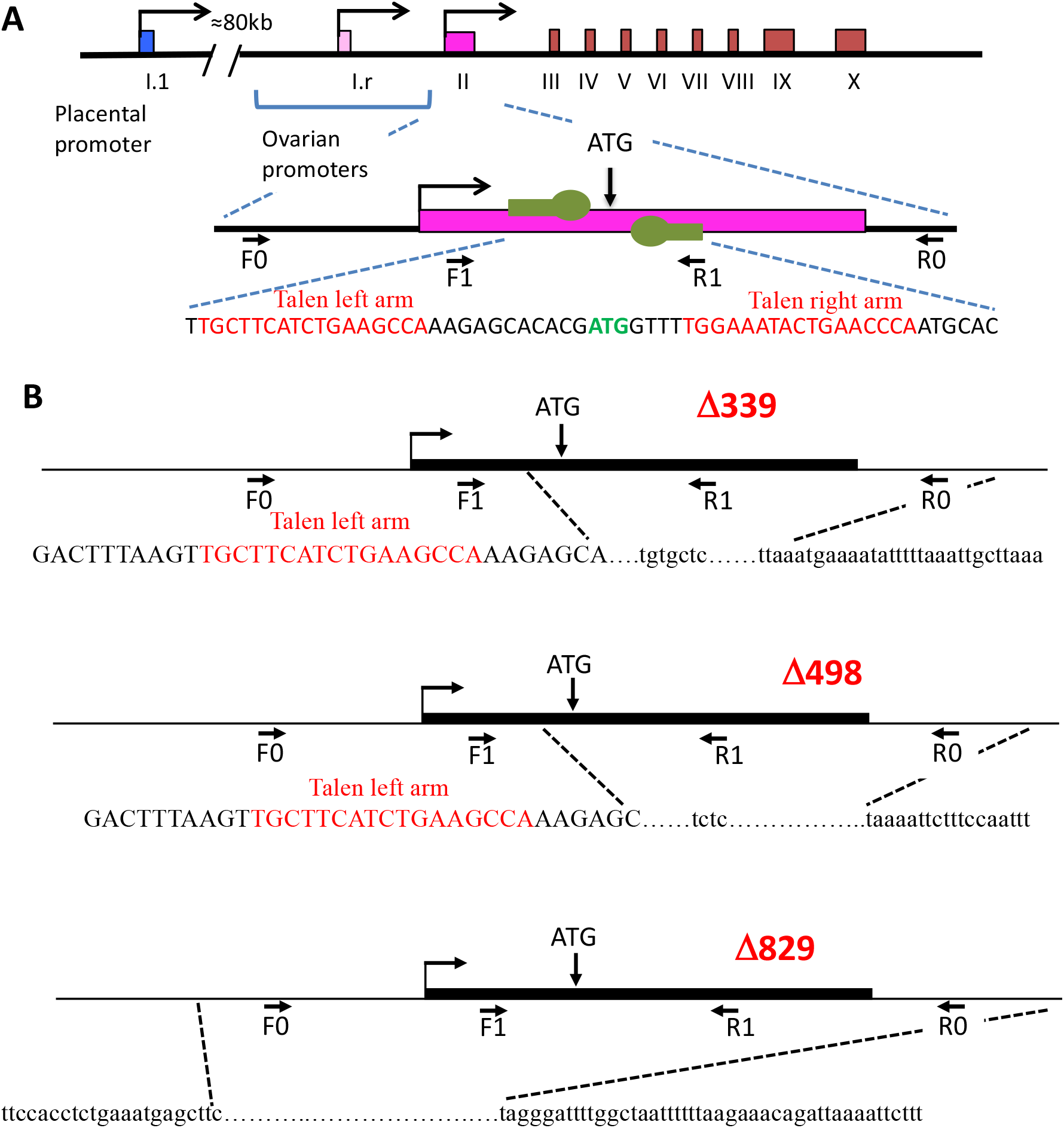
TALEN induced indel mutations in the rabbit *CYP19A1* gene. **(A)**: structure of the rabbit aromatase gene, with the various tissue specific promoters i.e. the I.1 placental promoter at around 80kb upstream of the transcription start point (tsp, marked by squared arrows) and the two proximal ovarian promoters I.r and II as previously described (Bouraïma *et al.* 2001). The enlargement shows the sequence of the exon II with the initiation site of translation (ATG in green letters). The two subunits (left and right arm) of the TALEN are shown as green symbols and the targeted sequence is written in red. The horizontal black arrows point the primers used for PCR detection of mutants in founders (F0/R0 set) and for routine qPCR genotyping (F1/R1 set). **(B):** sequences of the mutant alleles Δ339, Δ498 and Δ829. Capital letters report to exon sequence, lowercase letters to intron sequence. The deletion spans over 339 nucleotides (nt) in the Δ339 mutant allele (149 nt in the 3’ end of exon II and 190 nt of intron 2) and over 498 nt in the Δ498 mutant allele (148 nt in the 3’ end of exon II and 350 nt of intron 2), both with insertion of few nt at the repair position. In the Δ829 allele, the mutation consisted in the elimination of 829 nt encompassing 250 nt upstream of the beginning of exon II (transcription start site of the gene as regard to the ovarian promoter), the totality of the second exon (263 nt) and 316 nt in the second intron. All mutations suppressed the ATG codon and the splice donor site at the 3’ extremity of exon II.

Three mutant lines harboring each one a specific mutant allele were generated and studied. Homozygous (*ARO^-/-^*) rabbits were generated by crossing heterozygous (*ARO^+/-^*) rabbits. Mating was performed between rabbits within each line in order to avoid mixing the various mutant alleles.

Mutations are shown in Figure 1. A 339, 498 or 829 nucleotide-long (nt) deletion was observed in each of the alleles respectively named Δ339, Δ498 and Δ829. Briefly, mutations eliminated the translation initiation site (ATG codon). Thus, no protein was expected to be translated from any of these mutant alleles. Otherwise, the deletions theoretically should lead to the elimination of 48 amino acids in the N-terminus of the AROMATASE enzyme. As previously published (Kaur and Bose 2014), these amino acids encompass the signal anchor responsible for the translocation of the protein to the endoplasmic reticulum. It is thus expected that, if a protein is translated from the mutant mRNAs, it has a good chance of not being properly integrated into the endoplasmic reticulum bilayer, and therefore non-functional.

Finally, to confirm the absence of any active AROMATASE enzyme, we searched for estradiol by Gas Chromatography / Mass Spectrometry (GC/MS) in rabbit serum samples collected after birth and in fetal gonads. The concentrations of estradiol in sera from *ARO^-/-^* XX females after birth were under the limit of quantification (LOQ = 0,2pg/ml) whatever the age of the animals, when in wild type (*ARO^+/+^*) and heterozygous (*ARO^+/-^*) females the estradiol concentration increased progressively with the age of the animal (supplementary Figure 1). The concentration of testosterone was measured in the same samples. In *ARO^-/-^* sera, the values ranged in similar way than in WT or *ARO^+/-^* XX rabbits.

We measured also by GC/MS the steroid content of the gonads in fetuses aged of 28 days *post coitum* (d*pc*). The estradiol content was undetectable in gonads from all *ARO^-/-^* XX fetuses and in all XY gonads, when it was always detected, even sometimes in very low amounts, in XX gonads from *ARO^+/-^* and WT genotypes (supplementary figure 2). Testosterone was not detected in any XX gonads whatever their genotype at the *CYP19A1* locus. This shows that androgens were not over-synthesized nor accumulated in XX *ARO^-/-^* gonads. Finally, these data also show that neither estradiol nor testosterone from maternal or placental origin was detected in the 28d*pc* fetal gonads.

### XY and XX *ARO^+/-^* rabbits were fertile, with normal genital tracts and normal gonads morphology

In the three mutant lines, all *ARO^+/-^* and XY *ARO^-/-^* rabbits were viable until adults and did not suffer apparently from any metabolic or postural disease at least until 2 years after birth. No apparent modification of external genitalia was visible. All were fertile and gave birth to progeny. The sexual behavior was not studied. The number of neonates per litter from heterozygous mating (**♀** *ARO^+/-^* x **♂** *ARO^+/-^*) was similar to that of wild type mating (**♀** WT x **♂** WT) (supplementary Figure 3). The percentage of mating without pregnancy was not different: 37,9% (11 null per 29 mating) in WT x WT mating and 35,3% (23 null per 65 mating) in heterozygous x heterozygous mating. All this suggests that the fertility of heterozygous animals was not affected.

At puberty (5-6 months after birth), genital tracts and gonads were not different in heterozygous and wild type XX rabbits (supplementary Figure 4). Since all measured parameters did not differ in heterozygous and wild type females (similar estradiol and testosterone serum levels, similar prolificacy, similar overall structure of genital tracts and ovaries), we used ovary samples indifferently from wild type or heterozygous females as reference ones. Nevertheless, the genotype has always been distinctly reported in the legends of all figures.

### XX *ARO^-/-^* rabbits developed small female genital tracts, ovaries with almost no follicular reserve and very few antral follicles

All XX *ARO^-/-^* rabbits were viable without any metabolic or postural disease at least until 1.5 years after birth. No apparent modification of external genitalia was visible. The sexual behavior of XX *ARO^-/-^* rabbits was not studied.

Obvious modifications of the genital tracts and ovaries were observed in XX *ARO^-/-^* rabbits from the three lines with similar characteristics. After puberty, a typical phenotype was observed, with under-developed uterus horns, oviducts and pavilions resembling those of neonatal females. Their size was clearly smaller than in wild type or heterozygous females (supplementary Figure 4). In all three lines, XX *ARO^-/-^* gonads developed as ovaries (Figure 2, supplementary Figure 5) but with a very small size. As the phenotype was identical in Δ339, Δ498, and Δ829 lines, it is hardly attributable to any off-target alteration of the genome. Ovaries contained almost no primordial follicles, scarce primary and secondary follicles, and were filled with numerous remnants of atretic follicles. The abundant follicle reserve composed of primordial follicles observed at the cortex of wild type ovaries (Figure 2A and B) was absent in *ARO^-/-^* gonad mutants (Figure 2C and D). Nevertheless, some follicles developed to reach the large antrum stage in *ARO^-/-^* ovaries, showing granulosa and theca cell layers, oocyte and cumulus. The number of large antrum follicles was high in some females (see the ovary of the Δ829 female in supplementary Figure 5) possibly reflecting the surge of gonadropins that occurs at early puberty (around 5-6 months in the rabbit). Numerous Call-Exner bodies (spherical deposits of basal lamina matrix surrounded by granulosa cells, observed in rabbits and some mammals) were observed in the granulosa of growing follicles in *ARO^-/-^* ovaries as classically observed in wild type ones (Lee *et al.* 1996). The number of pyknotic cells or of cells with dense colored nuclei was low.

**Figure 2:**
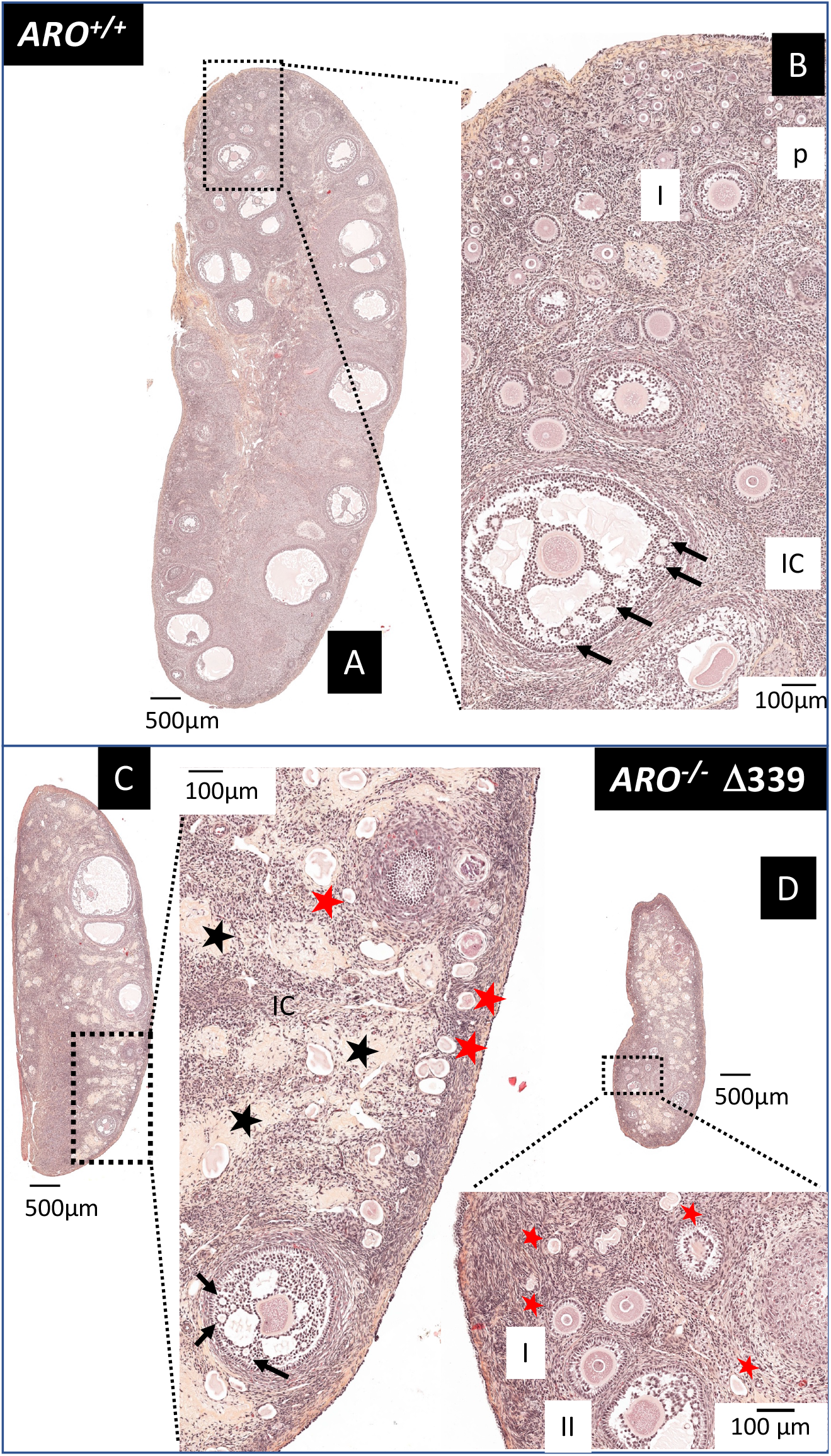
ovaries of adult *ARO^-/-^* rabbits were small with few follicles and filled with collagen rich connective tissue. Ovaries from wild type female rabbits (*ARO^+/+^,* A and B) and from the mutant *ARO^-/-^* line Δ339 (C and D) were fixed in PAF then HES colored. Females were 5,5 months old. Two distinct sections of the same ovary are presented for the mutant *ARO^-/-^* to show the follicles that were otherwise not visible. Enlarged zones point the different types of follicles. Black arrows = Call-Exner bodies. Black stars = accumulation of fibrous tissue. Red stars = remnants of degenerated follicles. “p” = primordial follicle; I = primary follicle; II = secondary follicle; IC = interstitial cells. The phenotype of the *ARO^-/-^* ovary was similar in the other lines Δ498 and Δ829 (supplementary Figure 5).

The expression of the marker genes *RSPO2* and *FOXL2* specific respectively for the oocyte and for the granulosa cells in differentiating follicles was studied by *in situ* hybridization (ISH, Figure 3). In the *ARO^-/-^* ovary, the rare detected oocytes were positive for the *RSPO2* labelling (Figure 3A), and granulosa cells were positive for *FOXL2* (Figure 3B). In wild type ovaries, AROMATASE was immuno-detected in a small number of follicles with antrum, thus characterizing the pre-ovulatory ones (Figure 3C). The Anti Müllerian Hormone (AMH) was immuno-detected in granulosa cells of many follicles in wild type ovaries except those positive for AROMATASE. In *ARO^-/-^* ovaries, granulosa cells of all follicles were AROMATASE negative and AMH positive (Figure 3D). However, the AMH serum level was lower in *ARO^-/-^* females, reflecting the small number of follicles and/or their poor synthesis capacity.

**Figure 3:**
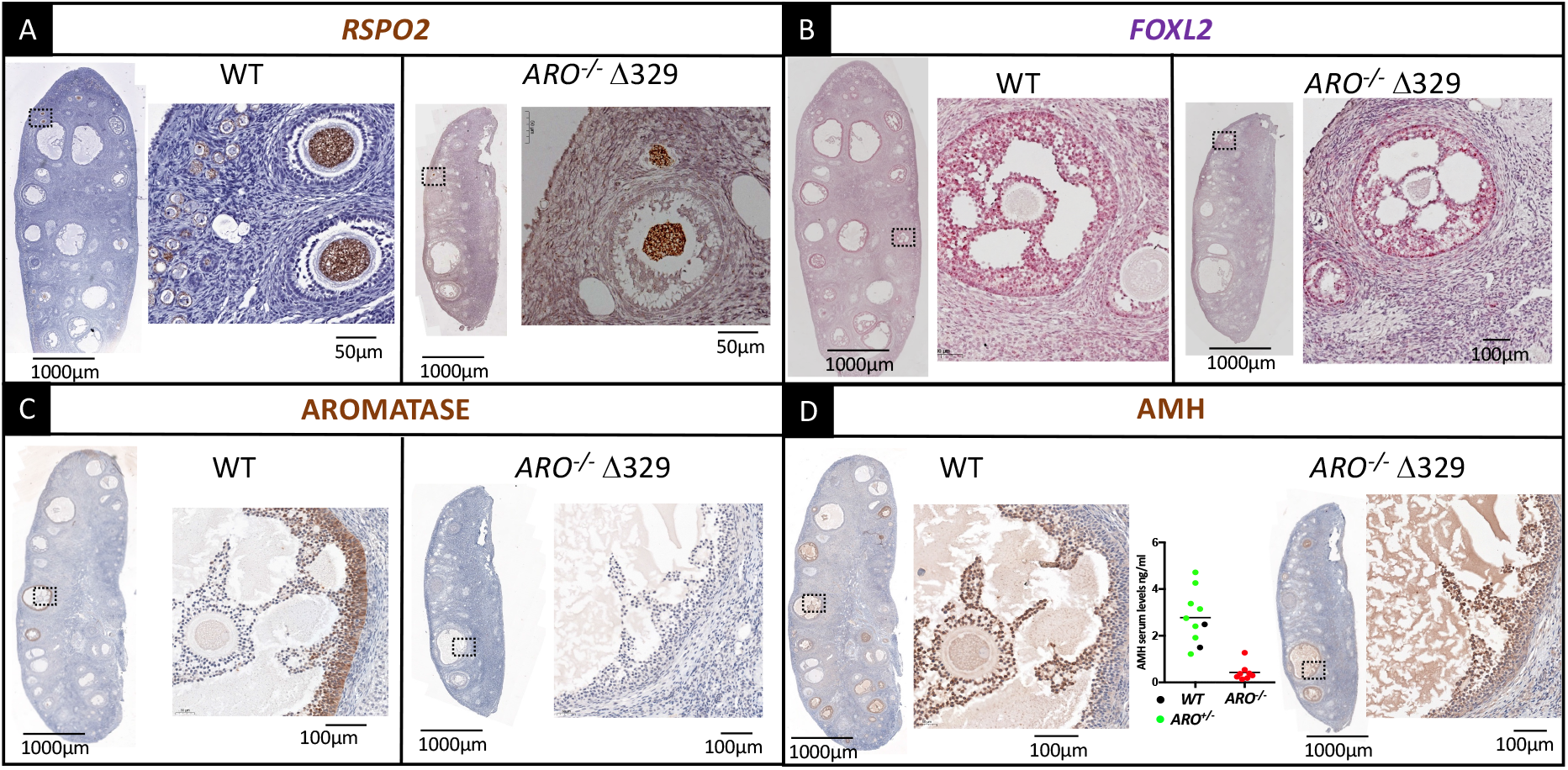
*RSPO2, FOXL2* and AMH were detected in wild type and ARO^-/-^ adult ovaries. In Situ Hybridization to localize mRNA of *RSPO2* and *FOXL2* genes and immunodetection of AROMATASE and AMH in adult ovaries. Ovaries were collected in wild type (WT) or *ARO^-/-^* Δ329 females aged 5-6 months. The expression of *RSPO2* (A) and *FOXL2* (B) genes was detected by ISH probes. AROMATASE (C) and AMH (D) were detected by immunohistochemistry on two adjacent sections. The *RSPO2* and *FOXL2* probes labelled respectively the cytoplasm of oocytes (brown dots) and of granulosa cells (red dots) from all follicles, from primordial to antrum stages in wild type and *ARO^-/-^* rabbits as well. Some thecal cells were also positive for the *FOXL2* labelling in both genotypes. Mural granulosa cells from pre-ovulatory follicles were positively labelled by the anti-AROMATASE antibody in wild type ovaries (brown colored); no cells were positive in *ARO^-/-^* ovaries. The anti-AMH antibody labelled most granulosa cells of growing follicles in wild type ovaries (brown colored), excepted those positive for AROMATASE. In *ARO^-/-^* ovaries, all follicles with antrum were positively labelled. Similar staining were observed in females from the two other strains. The graph represents the AMH serum level in *WT* (black points), *ARO^+/-^* (green points) and *ARO^-/-^* (red points) females.

Intense modifications affected the medulla of the ovary. In wild type rabbits, the medulla consisted mostly in interstitial tissue composed of cells with large clear cytoplasm and round regular shape nuclei (Figure 2B); besides, in *ARO^-/-^* ovary, the interstitial tissue appeared as a mixture of cells with heterogeneous shape and angular nuclei. More, several areas filled with fibrous material rich in collagen and devoid of cells were detected (Figure 2C and D; supplementary Figure 5).

Some of the heterozygous (*ARO*^+/-^) and homozygous (*ARO*^-/-^) 10 months old XX rabbits were hormonally treated to induce ovulation, mated and sacrificed to collect gonads for histological analysis. Ovulation rupture points were observed in ovaries from *ARO*^+/-^ rabbits, but not in *ARO^-/-^* ovaries (supplementary Figure 6). While several primordial, primary, pre-antral and antral follicles were detected at the cortical periphery of ovaries in *ARO*^+/-^ rabbits after super-ovulation treatment, no primordial or primary follicles were seen in the treated *ARO^-/-^* ovarian cortex. Nevertheless, some large antrum type follicles were observed in *ARO^-/-^* ovaries, with developed granulosa, theca cells, and oocyte.

### The mutation impacted the XX gonad from the first stages of their differentiation

Our goal was to search when and how ovaries were impacted by the lack of functional AROMATASE enzyme and of estradiol during fetal life. In the rabbit species, pregnancy spans for 31 days. Transition from an indeterminate gonad to an ovary or testis occurs at around 16-18 d*pc*. The peak of *CYP19A1* gene expression starts at 18 d*pc* in the ovary, concomitantly with the increase of expression of the *FOXL2, ESR1, WNT4* and *RSPO1* genes (supplementary figure 7 and (Daniel-Carlier *et al.* 2013). The *RSPO1* and *WNT4* genes are major components of the β-catenin signaling already reported as mandatory for ovarian early differentiation in mice and humans (Vainio *et al.* 1999; Parma *et al.* 2006; Chassot *et al.* 2012). The *ESR1* and *ESR2* genes are those coding for the α- and β-estrogen receptors respectively. The *CYP19A1* and the *ESR1* genes were expressed by cells of the surface epithelium and those forming invaginated structures issued from this epithelium (20 d*pc*, supplementary figure 7, ISH probes labelling). On the other hand, the *FOXL2* gene was mainly expressed in somatic cells underneath the surface epithelium and those surrounding the invaginated structures. The expression of the *ESR2* gene coding for the estrogen receptor beta was low at these stages compared with the levels determined later at birth (34 d*pc*). As expected, a faint labelling was observed by ISH labelling (supplementary Figure 7).

Therefore, the present study analyzed the fetal gonads from 18 d*pc* onwards. The gonads from all XX fetuses, *ARO^-/-^* and *ARO^+/-^,* were differentiated as ovaries with the same morphology except a thinner coelomic epithelium in *ARO^-/-^* gonads (Figure 4A). Regardless of the developmental stages, XX *ARO^-/-^* gonads showed no evidence of testicular appearance (see for comparison the 18 d*pc* testis in the supplementary Figure 8). A similar density of OCT4 positive germ cells was detected in *ARO^-/-^* and *ARO^+/-^* gonads (similar number of positive cells per surface units, Figure 4B). Mitotic activity was detected by KI67 immunostaining (Figure 4C). KI67 positive cells were mainly present in the ovarian surface facing the abdominal cavity and in invaginated structures below in both *ARO^+/-^* and *ARO^-/-^* ovaries. Interestingly, the KI67-positive ovarian surface appeared thinner in KO gonads than in the *ARO^+/-^* controls (Figure 4C).

**Figure 4:**
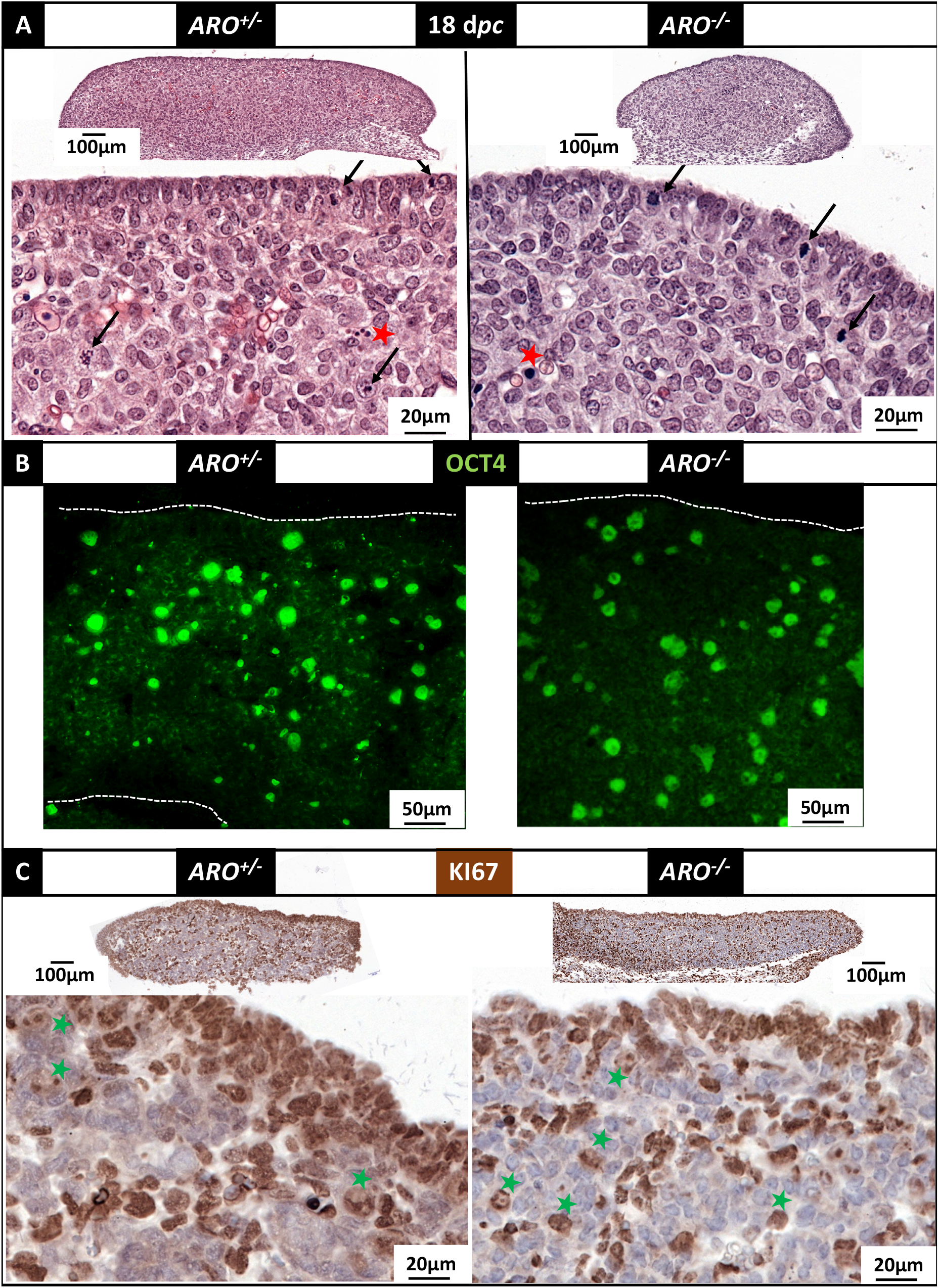
morphology of ovaries from *ARO^+/-^* and *ARO^-/-^* 18 *dpc* old fetuses and detection of OCT4 and KI67. Gonads were fixed in Bouin’s then HES colored (A) or fixed in PAF then treated for antibody labelling (B and C). **(A): HES coloration**. The surface epithelium appeared as a continuous layer of epithelial cells. Red stars point dense nuclei that could correspond to pyknotic nuclei. Arrows point figures of mitoses. **(B): detection of OCT4 positive cells.** The immune-labelling was visualized using a fluorescent secondary antibody. Nuclei of positive cells (large round green-labelled) were dispersed through the gonad in both *ARO*^+/-^ and *ARO*^-/-^ ovaries. **(C): detection of KI67 positive cells.** The immune-labelling was visualized using a peroxidase coupled secondary antibody. Positive cells with brown colored nuclei were found at the surface epithelium, and in rows of cells delimiting cell clusters. Green stars point brown light-colored large nuclei, possibly corresponding to germ cells.

From 20 d*pc* onwards, more important alterations progressively appeared in *ARO^-/-^* ovaries. The cortex was thinner, composed of a single layer of epithelial cells, connected to small size nascent ovigerous cords, by opposition to the thick cortex observed in *ARO*^+/-^ ovaries, connected to numerous large size ovigerous cords (Figure 5A and 5D). A discontinuous layer of cells with small and dense colored nuclei was observed underneath the surface epithelium, forming at 22 d*pc* (Figure 5D) and later on (34 d*pc*, Figure 9) a loose connective tissue between the thin cortex and the inside of the ovary.

**Figure 5:**
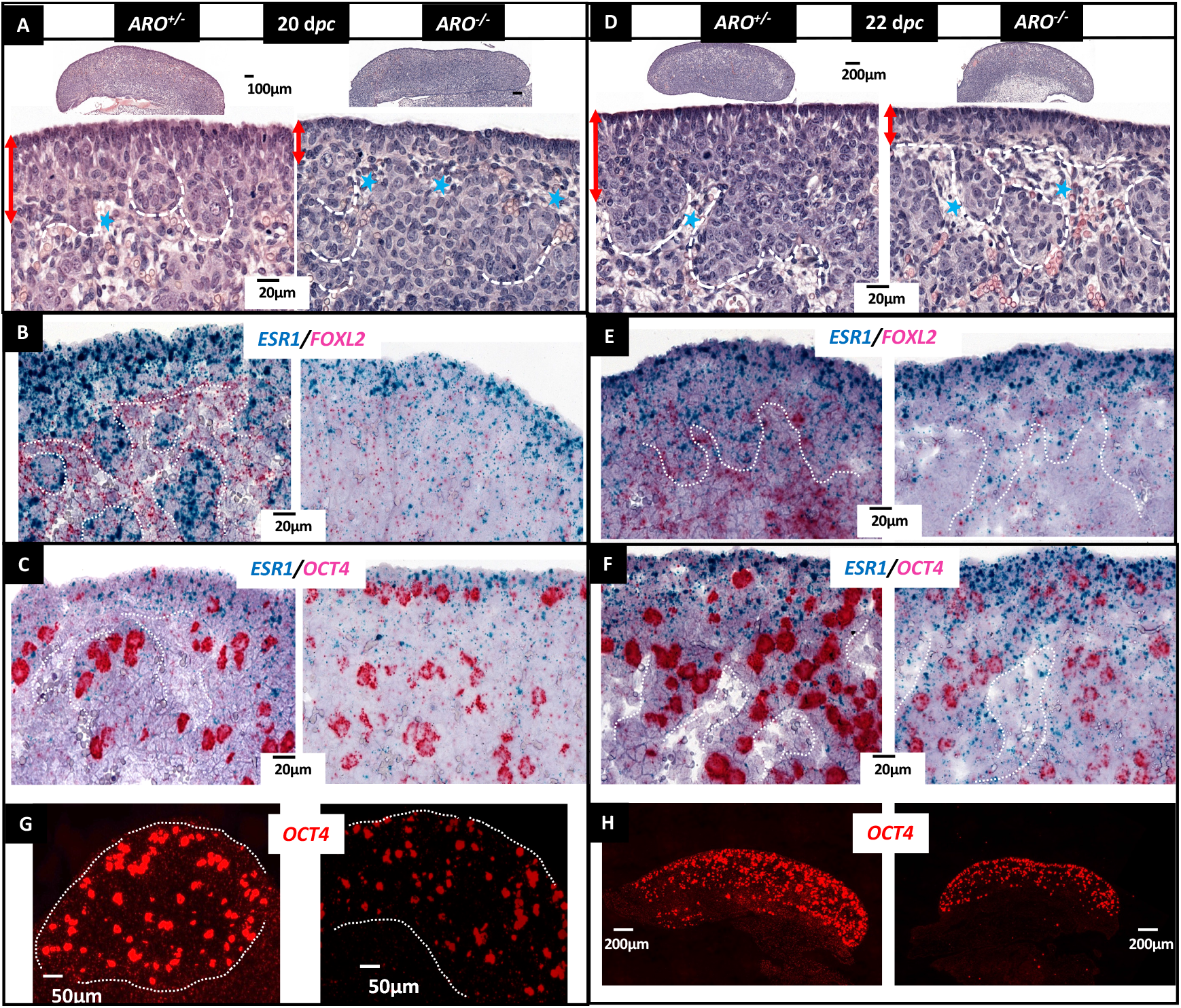
gradual modifications of *ARO^-/-^* ovaries at 20 and 22 d*pc*. Ovigerous nests and cords are underlined by a dotted white line. The red double arrow points the thickness of the nascent cortex **(A, D)**. Blue stars point the connective tissue. *In situ* hybridization was performed to visualize simultaneously two RNA targets to show the *ESR1* (blue points) and *FOXL2* (fast red points) RNAs **(B, E)**, and the *ESR1* (blue) and *OCT4* (fast red) RNAs **(C, F)**. The lower panel is the fluorescence observation of the fast red *OCT4* ISH labelling to show the overall density of the *OCT4* positive cells (**G, H**).

The cells of the surface epithelium expressed the *ESR1* gene in both genotypes (Figures 5B and 5E). The intensity of the ISH *ESR1* labelling varied greatly at the surface of the *ARO^+/-^* ovaries as shown in figure 5 (compare 5B, E, C and F, which are different sections from the same gonad). Nevertheless, in all treated sections, clusters of labelled *ESR1* cells connected to the surface epithelium were easily observed. Besides, in *ARO^-/-^* ovaries, labelled *ESR1* cells were mainly dispersed in the inner of the gonad. Similarly, the *FOXL2* ISH probe labelled evenly dispersed cells in the inner of the *ARO^-/-^* gonad, which differed from the strong *FOXL2* labelled cell cords of the *ARO^+/-^* ovary. Numerous *OCT4* positive germ cells were found in the surface epithelium, interestingly in close contact with somatic cells expressing the *ESR1* gene. The hybridization signals were lighter in *ARO^-/-^* germ cells, thus in favor of a modification of their pluripotency/differentiated stage. As at 18d*pc*, their density was apparently not modified (Figures 5C and 5F-H). However, considering that the size of the ovary was smaller in 20 d*pc* KO fetuses, it is likely that the total number of germ cells was lower in KO than WT ovaries.

KI67 positive cells were overall detected in both types of ovaries at 22 d*pc* (Figure 6A) but there was still a clear difference of mitotic activity in the coelomic epithelium between *ARO^+/-^* and *ARO^-/-^* ovaries. Indeed, while many KI67 positive cell layers were detected in the coelomic epithelium of *ARO^+/-^* gonads, only one layer was visible in the *ARO^-/-^* one. The gH2AX-antibody, marker of double strand breaks of the DNA, labelled mostly germ cells in both types of ovaries as suggested by their large round nuclei (Figure 6B). The density of gH2AX positive cells (probably apoptotic cells) was apparently not different in *ARO^+/-^* and *ARO^-/-^* ovaries.

**Figure 6:**
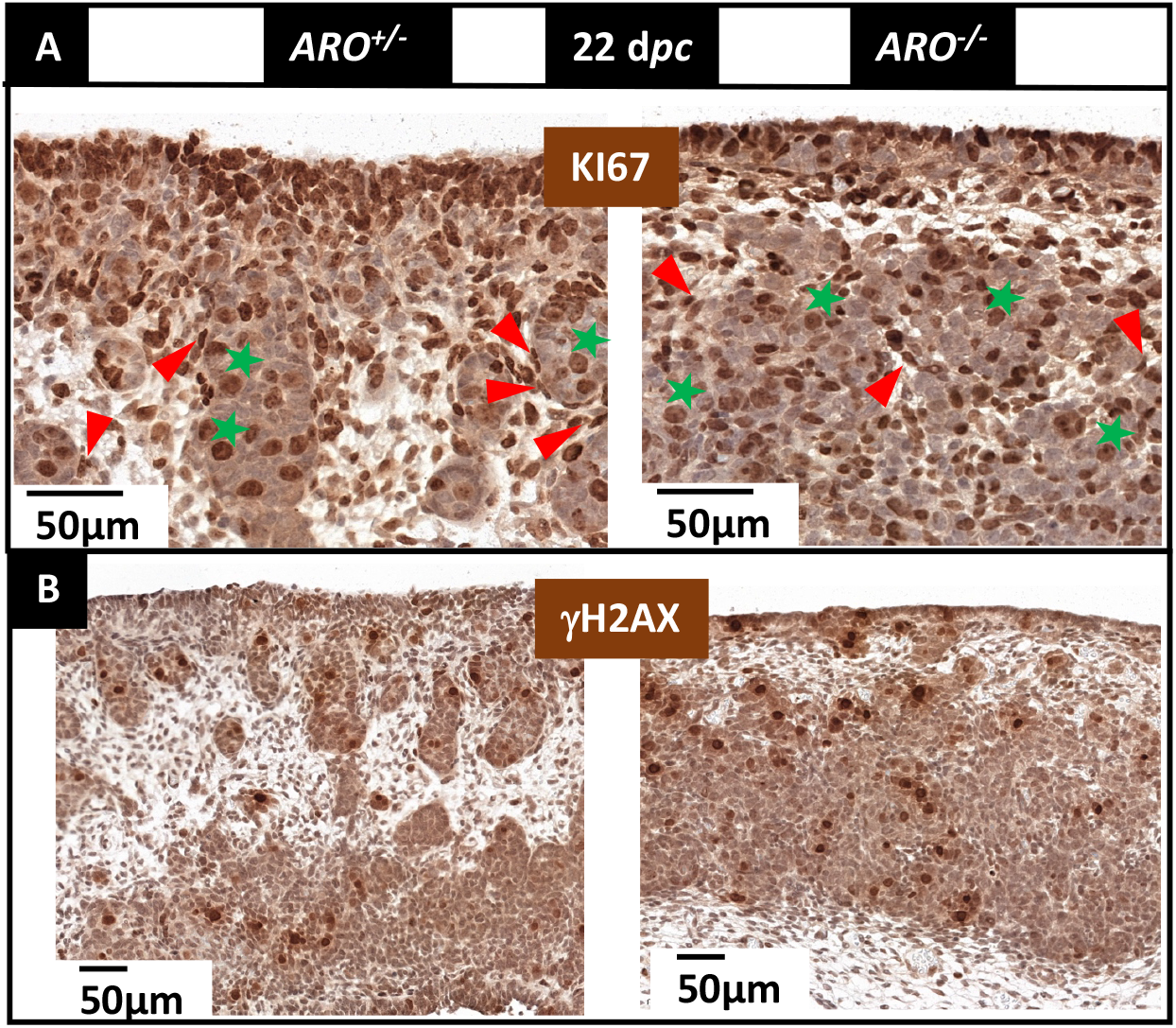
mitotic activity and DNA fragmentation at 22 d*pc*. The marker of mitotic activity KI67 **(A)**, and that of double strand breaks gH2AX **(B)** were localized by immunohistochemistry in sections from PAF fixed ovaries of *ARO^+/-^* and *ARO^-/-^* fetuses at 22 d*pc*. The KI67 antibody labelled nuclei of most cells of the coelomic epithelium, and cells inside the ovary in both genotypes. Cells with flat nuclei KI67 positive (red arrowheads) surrounded from place to place the ovigerous cords. Cells with large round nuclei positive for KI67 labelling (probably germ cells) were found mainly at the inside of the cords (green stars). The gH2AX antibody labelled large round shape nuclei corresponding probably to germ cells. Note the disconnection between the coelomic epithelium and the new forming ovigerous cords in *ARO^-/-^* ovaries.

The relative expression levels of several key marker genes were determined on total RNAs extracted from the whole gonads (Figure 7). Expressions of *OCT4, FOXL2* and *RSPO1* genes were reduced in *ARO^-/-^* ovaries, especially at 22d*pc*, reflecting a reduction of the number of cells or an alteration of their differentiation, or both. Interestingly, that of *WNT4* was significantly higher at 22 d*pc* in KO ovaries. The ratios *RSPO1/FOXL2* and *WNT4FOXL2* were calculated to analyze the relative variations of these somatic cell specific genes. Ratios were similar at 20d*pc*. However, two days later, at 22d*pc*, both were significantly higher in KO ovaries. At last, this shows that the two genes *RSPO1* and *WNT4* were relatively more expressed in KO ovaries at 22d*pc*, and suggests that the β-catenin signaling was stimulated in KO ovaries. The expression level of the *DDX4* gene, known to be its lowest at this ovarian developmental stage (Daniel-Carlier *et al.* 2013), did not differ between WT and *ARO*^-/-^. It increased between 20 and 22 d*pc*, which reflects the early differentiation of germ cells from the pluripotent stage. The *SOX9* gene expression level was 5- to 10 times lower in the ovaries from all genotypes than in testes at the same age. The SOX9 transcription factor was not detected by immunohistology either in wild type or in *ARO*^-/-^ ovaries, while a clear labeling was observed in the fetal testis at 22 d*pc* (supplementary Figure 8). This reinforces the demonstration that *ARO*^-/-^ ovaries did not show any sex-reversal phenotype. The expression levels of the *ESR1* and *ESR2* genes did not change significantly.

**Figure 7:**
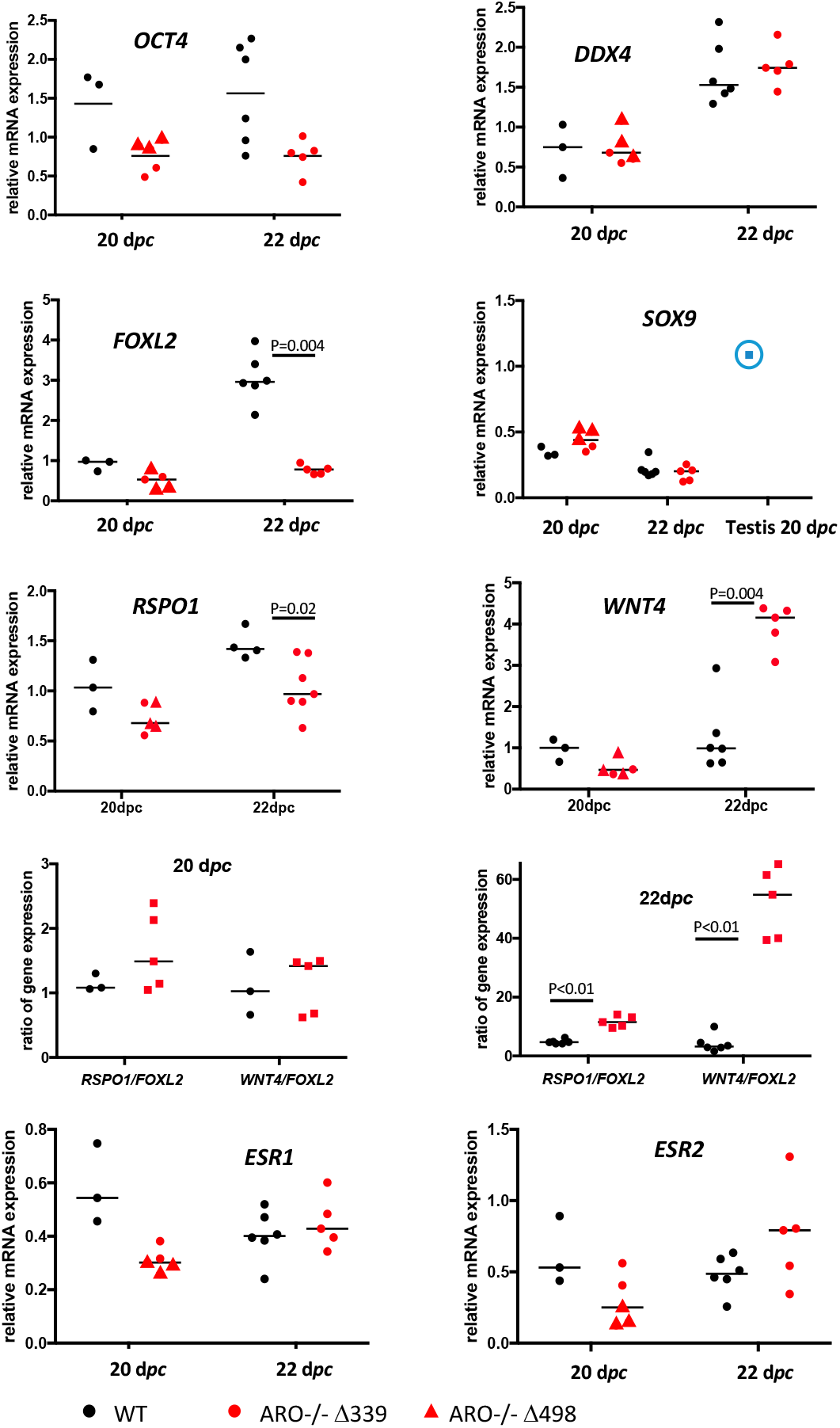
expression of somatic and germ cell specific marker genes in 20 and 22 *dpc* ovaries from WT and *ARO^-/-^* fetuses. The diagrams show the relative mRNA expression levels. Each point represents one RNA sample extracted from both gonads of one animal at the indicated developmental stage. The same RNA samples were analyzed in the diagrams. Red triangles refer to mutant from Δ339 line, and red point from Δ829 line. No samples from line Δ498 were analyzed. The blue point in the *SOX9* diagram shows the *SOX9* mRNA level measured in the two testes of one 20 d*pc* WT fetus. Horizontal bars represent the medians. *DDX4* and *ESR2* gene expression levels were very low, but significantly measured. The 20d*pc* and 22d*pc* diagrams represent the ratios of gene expression *RSPO1/FOXL2* and *WNT4/FOXL2* at 20 and 22 d*pc* respectively. Values are those of the mRNA expressions given in the above diagrams.

Thus, from 20 d*pc* onwards, the differentiation of the *ARO^-/-^* gonad was profoundly altered, with a decrease of proliferation of the somatic cells of the coelomic epithelium, a reduction of the number of germ cells, and alterations of the differentiation of all cell types. Finally this leads to a clearly underdeveloped small organ with small ovigerous nests, the spread of connective tissue underneath the surface epithelium, an abnormal distribution and differentiation of the *FOXL2* positive cells and alterations of expression of genes involved in ovary differentiation, such as the β-catenin signaling.

### Impact of estrogens deprivation on germ cell meiosis and follicular differentiation

In the rabbit ovary, the first signs of meiosis are histologically detected at 30-31 d*pc,* i.e. around birth. Later on, typical meiotic germ cells (prophase I) are observed for less than 2 weeks. After meiosis arrest at diplotene stage, follicles begin to differentiate. The peak of expression of the *STRA8* gene that starts at 24-28 d*pc* (Daniel-Carlier *et al.* 2013) marks the commitment of germ cells to meiosis. Thus, we sought to analyze at 28 d*pc* (meiosis commitment), 34 d*pc* (beginning of meiosis) and few days or weeks after birth (folliculogenesis) whether initiation and arrest of meiosis I and early follicular formation were impacted in *ARO^-/-^* ovaries.

In wild type and *ARO^-/-^* 28 d*pc* ovaries, numerous germ cells were positively labelled by the *OCT4, DDX4* and *STRA8* ISH specific probes (Figures 8B, 8C, 8D). The *STRA8* labelling proves the overall commitment of germ cells in meiosis in WT and *ARO^-/-^* ovaries. However, in these later, the number of positive germ cells for the three probes was low. More, they were clustered in small ovigerous-like structures, while they were all observed in large ovigerous cords in wild type ovaries. Positive *FOXL2* labelled cells were also much less numerous in *ARO^-/-^* ovaries, as shown by the few numbers of hybridization labels. Nevertheless, some of these *FOXL2* positive cells harbored a flattened shape and surrounded the small ovigerous-like structures embedding germ cells, as in wild type ovaries (Figure 8E).

**Figure 8:**
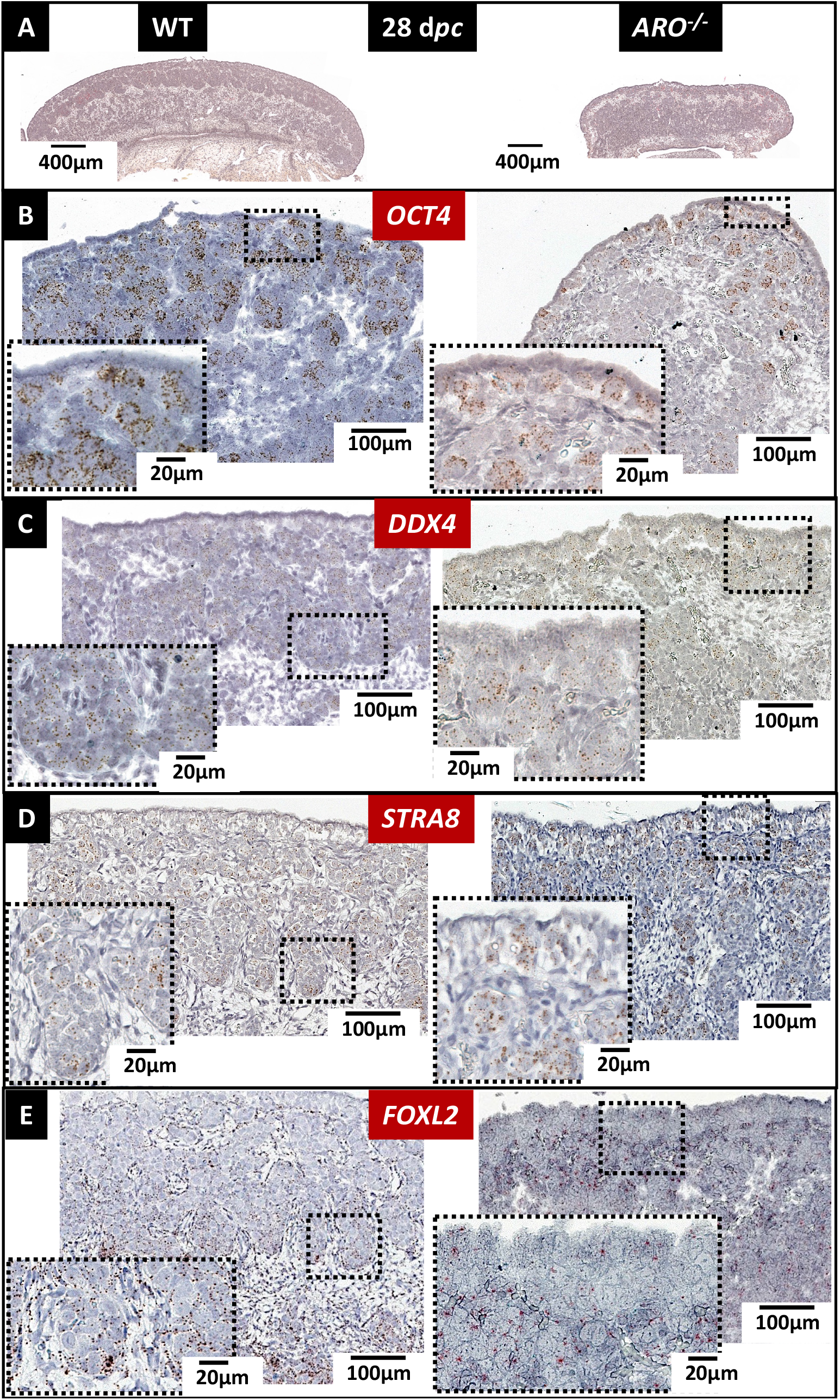
germ cells were committed to meiosis in WT and *ARO^-/-^* ovaries from 28 d*pc* old fetuses. Ovaries were fixed in PAF, then HES colored or analyzed by ISH. **(A): morphology of the ovaries**. The pictures show the difference of the overall size of the gonad and the different thickness of the cortex. The *OCT4* **(B)** and *DDX4* **(C)** ISH probes labelled respectively pluripotent germ cells and differentiating ones. The *STRA8* ISH probe **(D)** labelled germ cells committed to meiosis. The *FOXL2* ISH probe **(E)** labelled the differentiating somatic cells. Note that the latter were mainly distributed around and inside the ovigerous cords in the WT ovaries, and in contrast were dispersed throughout the gonad in the *ARO^-/-^* rabbit. As in all ISH pictures, positive labeling appears as colored dots.

**Figure 9:**
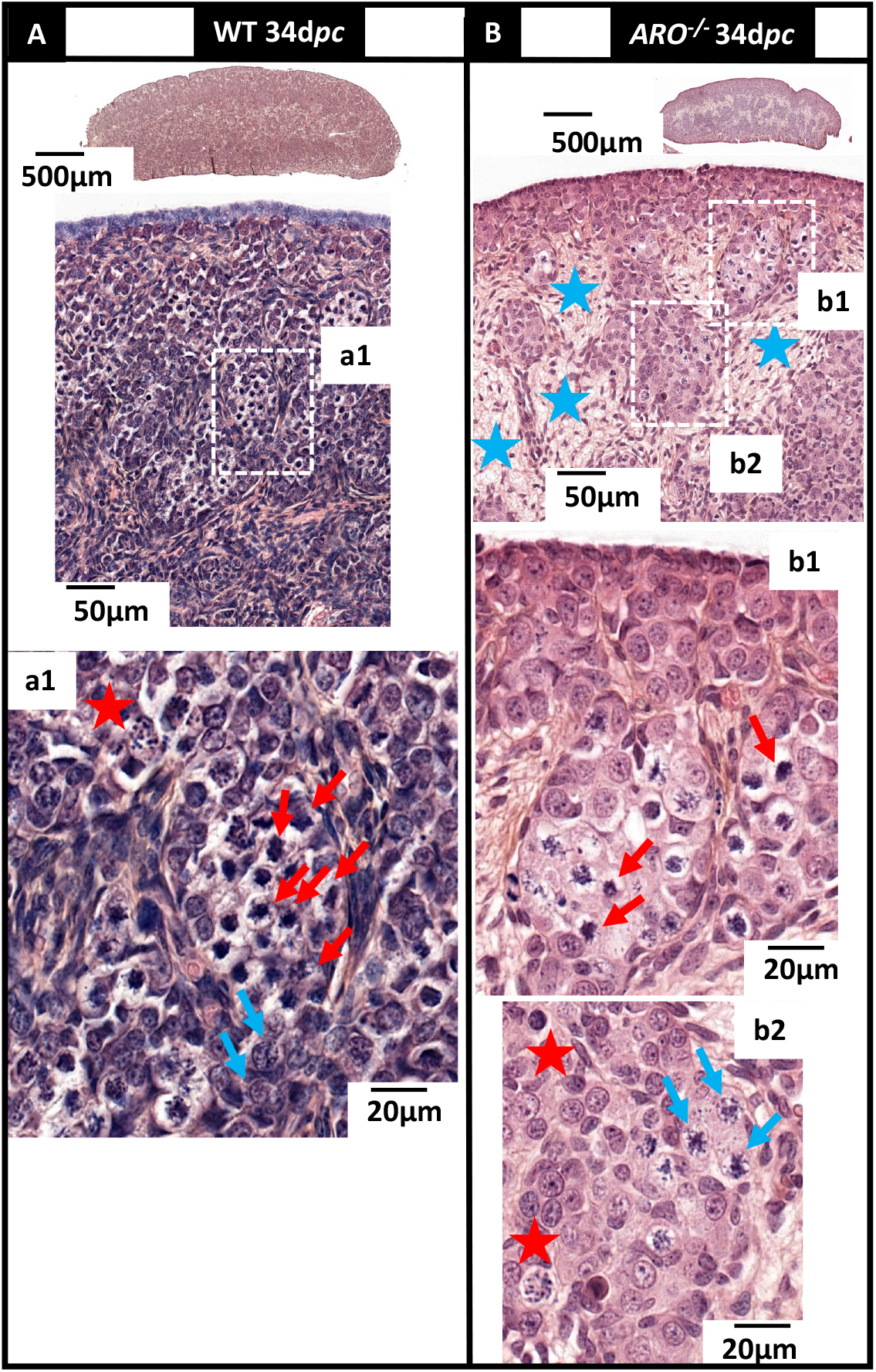
meiosis in WT and *ARO^-/-^* ovaries. Ovaries collected at 34 d*pc* (3-4 days after birth) were fixed in Bouin’s then HES colored. The stages of meiosis were identified as previously published (Peters *et al.* 1965). **(A):** In wild type ovaries, large ovigerous cords were filled with numerous germ cells in zygotene (a1, red arrows). Few pachytene stages were observed at the inner part of the cords (a1, blue arrows). **(B):** In *ARO^-/-^* ovaries, the rare ovigerous cords were filled with a few numbers of germ cells in zygotene (b1). As in wild type ovaries, some germ cells in pachytene were found at the inner part of the cords (b2). In both ovaries, aberrant pictures of cell nuclei were characterized by punctiform or condensed and dark colored chromatin (red stars). Blue stars show the connective tissue already observed mostly in *ARO^-/-^* ovaries.

At 34 d*pc*, in wild type ovaries, most ovigerous cords were filled with numerous zygotene germ cells, thus showing that meiosis had begun in a majority of germ cells (Figure 9A and a1). In contrast, in the *ARO^-/-^* ovaries, germ cells at zygotene were sparse (Figure 9B and b1, b2).

The differentiation of the *ARO^-/-^* ovaries was further altered as shown by the low staining observed with *DDX4, STRA8, SPO11* and *FOXL2* probes. The expression of all studied marker genes was reduced except that of the *WNT4* gene (Figure 10). The ratios of the *STRA8/DDX4* and *SPO11/DDX4* mRNA levels were lower in *ARO^-/-^* ovaries, suggesting that germ cells expressing *STRA8* and *SPO11* gene expression, and thus undergoing meiosis, were less abundant in *ARO^-/-^* ovaries. In contrast, the high ratio *WNT4/FOXL2* suggests as previously a relative activation of transcription of *WNT4* in somatic cells of KO ARO ovaries.

**Figure 10:**
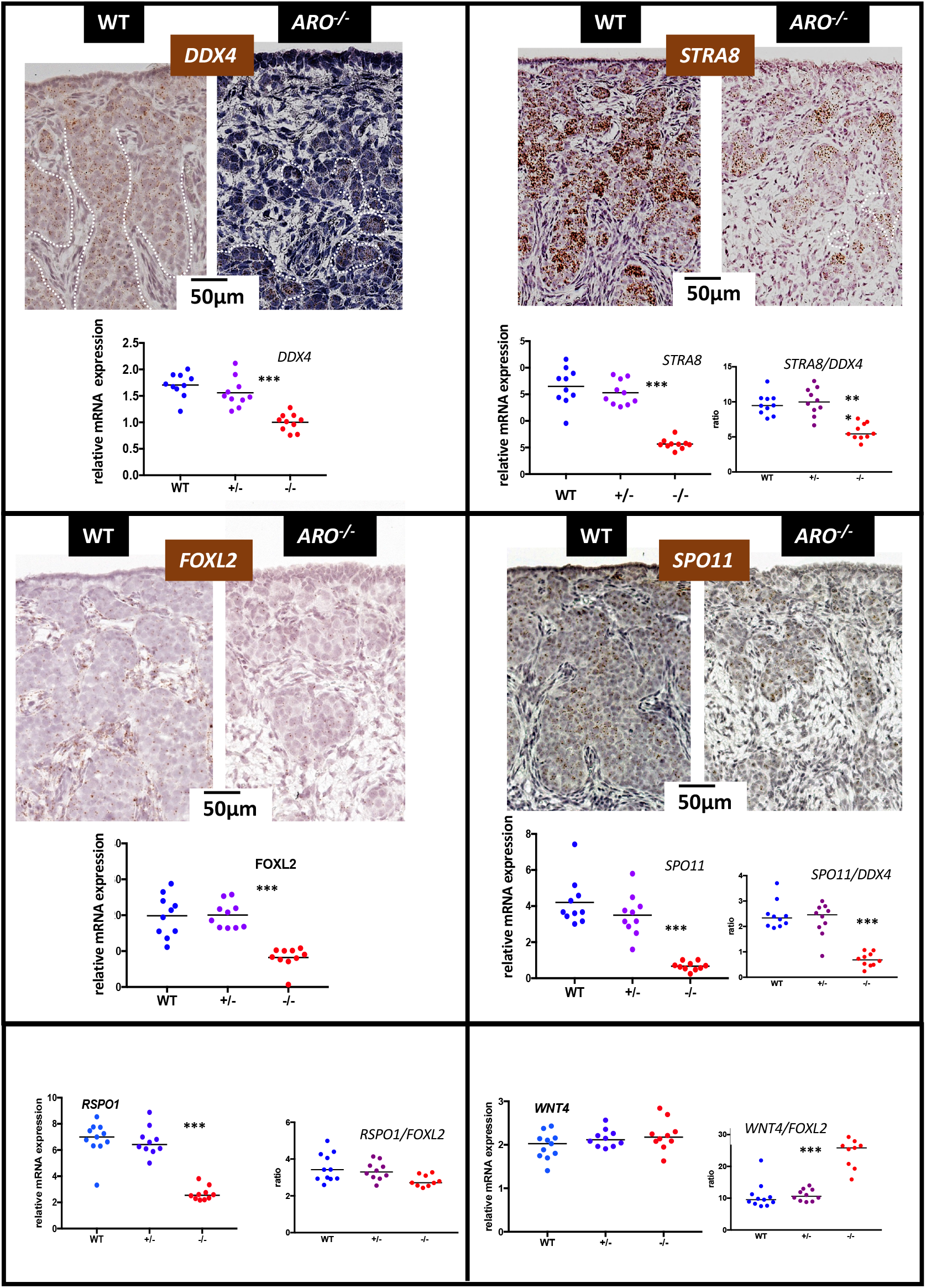
Marker gene expressions at meiosis. Germ cell marker genes (*DDX4, STRA8, SPO11*) and somatic cell marker ones (*FOXL2, RSPO1, WNT4*) were analyzed using ISH specific probes and quantitative PCR. In wild type ovaries, numerous germ cells positive for *DDX4, STRA8* and *SPO11* filled the ovigerous cords; pre-granulosa cells *FOXL2* positive were found around and inside the cords. In *ARO^-/-^* ovaries, germ cells positive for these same ISH probes were also found but much less numerous; the *FOXL2* labelling was very faint. The diagrams show the results of the quantitative analysis of gene expression. Values are relative expressions measured using total RNA extracted from the two gonads of each animal from wild type (WT), *ARO^+/-^* (+/-) *and ARO^-/-^* (-/-) rabbits; the same RNA samples were analyzed in the diagrams. Horizontal bars represent the medians. The expression of the germ cell marker genes *STRA8* and *SPO11* and that of the somatic marker ones *WNT4* and *RSPO1* were normalized respectively by *DDX4* and *FOXL2* gene expressions. Stars represent the statistical significance (p<0.01).

These differences intensified further (supplementary Figure 9). As meiosis progressed (10 days *post partum,* d*pp*), zygotene, pachytene and diplotene stages were simultaneously observed in wild type ovaries. At the interface between the cortex and the medulla, the large ovigerous cords broke down, forming numerous early primordial follicles encompassing diplotene stage germ cells surrounded by a ring of flat pre-granulosa cells. This was far to be observed in *ARO^-/-^* ovaries. Indeed, at 7 days after birth, almost no germ cells in zygotene or pachytene stages were detected; a small number of diplotene-like stage cells were nonetheless detected, some surrounded by a discontinuous ring of somatic cells that could be assumed to be pre-granulosa cells.

Later on at 16-19 d*pp*, the gonad size was approximately twice as small in *ARO^-/-^* rabbits, harboring a thin cortex with a small number of primordial follicles (Figure 11A). All ovigerous cords broke down both in wild type and *ARO^-/-^* ovaries and follicles were isolated. However, in *ARO^-/-^* ovaries, the ring of pre-granulosa cells surrounding the oocytes was frequently discontinuous and rare well-developed primordial follicles were detected. The *FOXL2* ISH labelling was very weak reflecting a low level of well-differentiated granulosa cells (Figure 11B). Nevertheless, oocytes of primordial follicles were positive for the *RSPO2* ISH specific probe in *ARO^-/-^* follicles, as in wild type ones (Figure 11C).

**Figure 11:**
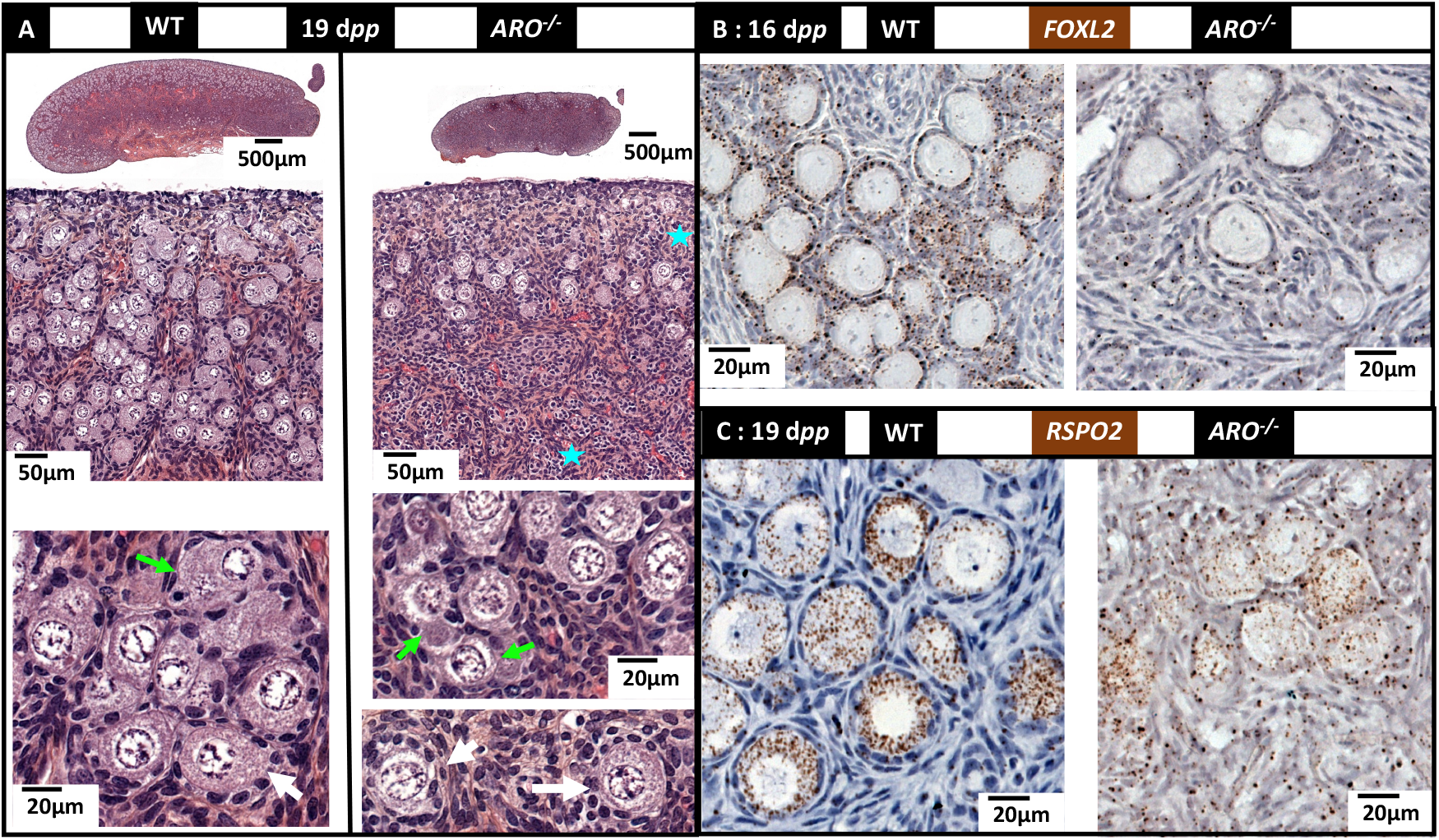
early follicle differentiation. Ovaries were collected at 16 or 19 d*pp* then treated with Bouin’s or PAF fixative and processed for HES staining or ISH labelling respectively. **(A): HES staining of 19 d*pp* sections**. Blue stars point the abnormal persistence of connective tissue, void of differentiating oocytes or follicles, at the outer part of the cortex and in the medulla of *ARO^-/-^* ovaries. The green arrows point some of the primordial follicles in formation with a discontinuous wall of granulosa cells. Early formed primordial follicles surrounded by a wall of regular shape granulosa cells are found in wild type and *ARO^-/-^* ovaries (white arrows). **(B) and (C): ISH labelling using *FOXL2* and *RSPO2* gene specific probes.** The *FOXL2* and *RSPO2* probes labelled granulosa cells and oocytes respectively in both wild type and *ARO^-/-^* ovaries. Note the huge decrease of *FOXL2* positive cells in *ARO^-/-^* ovaries compared to the WT control ones.

To conclude, in the *ARO^-/-^* ovaries, a small number of oocytes had differentiated after birth; these oocytes were surrounded by a ring of granulosa cells. Despite the probably abnormal differentiation of the latter (shown by the weak expression of the *FOXL2* gene), a small population of primordial follicles could differentiate. At the same time, the ovary was invaded by the connective tissue filled with fibers rich in collagen as previously observed from 20 d*pc* onwards; this latter was not replaced by the ovigerous nests as normally occurs in non-mutant ovaries.

## Discussion

The already long-standing demonstration of a peak of *CYP19A1* gene expression, AROMATASE activity and estradiol production by the fetal ovary in several mammal species between the stage of sex determination and the initiation of meiosis has raised questions on the role of locally produced estrogens on the fetal differentiation of this organ. The mouse model was for a long time the sole available model to study the function of genes *in vivo.* So, strains of *Cyp19a1* gene or *estrogen-receptor* genes KO mice have been created and a series of papers have reported their phenotype (Fisher *et al.* 1998; Couse *et al.* 1999; Britt *et al.* 2000; Britt *et al.* 2001; Findlay *et al.* 2001; Hamilton *et al.* 2014). All XX mice developed ovaries, but follicle differentiation was abnormal after birth. However, this model could not answer to our question since the mouse is one species where *Cyp19a1* gene expression is detected after meiosis initiation in the “late” fetal ovary (Dutta *et al.* 2014) and not before (Greco and Payne 1994). Thanks to the recent possibility to target genome modifications in numerous other species, we created a rabbit model harboring a mutation in the *CYP19A1* gene (*ARO^-/-^),* a species with a peak of *CYP19A1* gene expression in fetal ovary before meiosis. The *ARO^-/-^* rabbits were totally void of any estrogen production. Particularly, the amount of estradiol in gonads of all *ARO^-/-^* fetuses was undetectable, which eliminates the possibility of any compensatory mechanism through estradiol from maternal or placental origins. More, no testosterone was detected in the *ARO^-/-^* fetal gonads, thus eliminating any possibility of interference due to the local accumulation of androgens in response to the lack of androgen aromatization. The key results of this study regarding XX *ARO^-/-^* ovary phenotype were: 1) no XX sex reversal in *ARO^-/-^* rabbits, 2) a small size ovary from early fetal stages as the consequence of a low mitotic activity, 3) an abnormal differentiation of *FOXL2* positive cells, leading to the poor differentiation of ovigerous nests 4) a quasi-absence of follicular reserve, linked to the reduction of the number of meiotic germ cells and to early folliculogenesis failure.

### AROMATASE is not an ovarian determining enzyme in mammals

The question of the role of estrogens in sex determination in mammals was prompted by the finding that in goats, the *FOXL2* gene was a main actor for ovarian determination, since the *FOXL2* gene loss of function caused an ovary-to-testis sex reversal of the fetal gonad (Boulanger *et al.* 2014). Previously, FOXL2 was shown to be a crucial transcriptional activator of the *CYP19A1* gene in goats (Pannetier *et al.* 2006) and fishes (Govoroun *et al.* 2004; Pannetier *et al.* 2006; Guiguen *et al.* 2010), and possibly in chicken (Govoroun *et al.* 2004). Thus, the question was raised to know whether FOXL2 could act on sex determination through *CYP19A1* gene transcriptional activation. This question was all the more important that in non-mammalian vertebrate species, anti-aromatase drugs induced female to male sex-reversal (see (Vaillant *et al.* 2001) in the chicken and (Yin *et al.* 2017) in fish). The rabbit was possibly an interesting model since, as in goats, the surge of expression of the *FOXL2* gene in the fetal ovary occurs when starts the peak of *CYP19A1* gene expression at the gonadal sex determination (Diaz-Hernandez *et al.* 2008; Daniel-Carlier *et al.* 2013). The results of the present study demonstrate that estrogens play no role on the ovary/testis determinism of the rabbit gonad. A similar finding had already been described in humans, where several cases of total *CYP19A1* gene inactivation were reported (Akcurin *et al.* 2016; Zhu *et al.* 2016; Mazen *et al.* 2017; Unal *et al.* 2018; Praveen *et al.* 2020). In spite of numerous symptoms affecting the development of gonads, of external genitalia and of breast in the girl, with multiple disorders in energy and bone metabolisms, there was no sex reversal of the human ovary as regard to the genetic sex (Zhu *et al.* 2016). In contrast, opposite data were reported in the mouse species, describing an increase in the serum testosterone levels in *Cyp19a1^-/-^* females (Fisher *et al.* 1998) and the presence of seminiferous-like structures in the gonads of adult *Esr1/Esr2* double knock out females (Couse *et al.* 1999; Amano *et al.* 2017). However, this has been controversial and not found by all authors.

### Estrogens are involved in early ovarian differentiation

The fetal ovarian differentiation is known to be intimately linked to mechanisms that involve cellular interactions and cell migration, depending on autocrine and paracrine signaling pathways. These processes involve genes coding for the components of the extracellular matrix and cell-cell adhesion molecules. Estrogens have been reported as potent actors in these mechanisms (Juengel *et al.* 2002; Gentilini *et al.* 2007; Wilhelm *et al.* 2007; Zalewski *et al.* 2012; Hummitzsch *et al.* 2013). We have shown that the *CYP19A1* and the *ESR1* genes were expressed by the ovarian cortical cells and more specifically by cells located within the coelomic epithelium (Figure 5B). We can therefore hypothesize that in rabbit *ARO^-/-^* ovaries, the absence of estrogens can lead to a dramatic decrease of proliferative activity in the coelomic epithelium, evidenced by the KI67 staining (Figure 6A). The decrease of *FOXL2* and *RSPO1* expression from 20-22 d*pc* could thus be related to the reduction of somatic cell number but also to an alteration of their differentiation (Figure 7). Overall, it results in a reduced size ovary as detected shortly after gonadal sex determination.

A similar situation has already been described in chickens. Indeed, in this species, the experimental down-regulation of the epithelial *Esra* gene was sufficient to severely affect the ovarian cortex differentiation (Guioli *et al.* 2020). It was thus supposed that the natural absence of *Esra* expression in the cortex of the right ovary of the hen was at least partially responsible for the asymmetric degeneration of the right ovary (Guioli *et al.* 2014).

### Other mechanisms likely compensate for the lack of estrogen

Notably, without estrogens, the proliferation of ovarian cells seemed to be affected but not abolished since some nests were clearly visible in *ARO^-/-^* gonads at birth although under-developed. So, given that estrogens are involved in ovigerous nests formation, other factors are likely to play a role in this process as well. It has been shown in mice that the *Rspo1/Wnt4/β-catenin* pathway is involved in cell proliferation of the coelomic epithelium of both sexes during the gonadal switch (Chassot *et al.* 2012). This pathway could thus be responsible at least in part of the formation of the few ovigerous nests in *ARO^-/-^* rabbit ovaries. To support this hypothesis, there is the fact that the expression of the *WNT4* gene was significantly enhanced in the KO ovary (Figure 7). This led us to suppose that in the absence of estrogen, the gene pathways involving β-catenin signaling were overactivated to compensate partially for the weak mitotic activity and to counter the impairment of the differentiation of the gonad. Transcriptomics studies at the single cell level should help to identify the cellular trajectories and characterize such mechanisms.

### The number of germ cells was affected by the lack of estrogens

One of the main features of *ARO^-/-^* rabbit ovaries is the decrease in the number of germ cells, clearly visible from 20-22 d*pc* onwards. In mammals, germ cells colonize early the genital crests (Juengel *et al.* 2002; Wilhelm *et al.* 2007; Hummitzsch *et al.* 2013). In the rabbit species, it starts at 9 d*pc* and is over at around 18 d*pc* (Chretien 1966), when starts the *CYP19A1* gene expression. Thus, the germ cell colonization occurs independently of *CYP19A1* expression. Therefore, in the *ARO^-/-^* ovary, in spite of the absence of estradiol, the initial stock of germ cells was not altered.

In mammalian ovaries, once germ cells enter the genital ridges, they multiply intensely by mitosis until meiosis. Then, the number of germ cells does not increase any more. During fetal life in rabbits, the mitotic activity of germ cells, which peaks around 16-18 d*pc*, remains high up to 26 d*pc* (Chretien 1966). This coincides with the strong expression of the *CYP19A1* and *ESR1* genes (Supplementary Figure 7). Interestingly, the germ cells, labelled at this stage by the *OCT4* HIS probe, were co-localized in the surface epithelium with cells expressing the *CYP19A1* and *ESR1* genes. Thus, as in the case of somatic cell differentiation, it is likely that the estradiol produced by AROMATASE acts as paracrine regulator of the fate of germ cells. Numerous articles have shown that in mammals estradiol stimulates mitotic activity in the fetal ovary (see review in (Auersperg *et al.* 2001; Chou and Chen 2018)). Otherwise, the cell death that continuously occurs in the ovary (Hussein 2005; Myers *et al.* 2014) could explain the low number of germ cells. Finally, the low number of *OCT4* positive cells also could result from an early differentiation of germ cells which thus lose their pluripotency. The latter hypothesis is unlikely, since the expression of the *DDX4* gene (that reflects the differentiation of germ cells from pluripotency) is similar in KO and WT ovaries at 20-22dpc. Regarding cell death, we studied the number of germ cells positive for the gH2AX marker which indicates DNA fragmentation, and at this stage the dying germ cells. In *ARO^-/-^* ovaries, this number was similar to that in wild type ones. Thus, our data suggest that this is probably not germ cell death or early differentiation, but rather the lack of germ cell divisions that was responsible for the low number of germ cells in the early *ARO^-/-^* ovary before meiosis.

### Germ cells were committed in meiosis in spite of the lack of estrogens

The commitment of germ cells in meiosis marks the first step of their differentiation as oocytes. In the rabbit species, it occurs at the third part of pregnancy (at around 28 d*pc*), and is characterized by the surge of the *STRA8* gene expression in germ cells specifically. Interestingly, in the *ARO^-/-^* ovaries, numerous germ cells were committed to meiosis as shown by the significant surge of *STRA8* gene expression; more, the various steps of the meiotic prophase were distinguishable, and the meiosis specific maker gene *SPO11* was significantly expressed as well. Thus, it suggests that nor the estradiol neither the AROMATASE enzyme were necessary for meiotic induction and progression. As discussed previously, the number of germ cells possible to be committed in meiosis was reduced in *ARO^-/-^* ovaries. More, it should be pointed out that the ratio of gene expression *STRA8/DDX4* and *SPO11/DDX4* was low in *ARO^-/-^* ovaries, showing that an abnormal loss of germ cells occurred at commitment or during meiosis. Thus, although estrogens are not mandatory for meiosis commitment, the fact remains that they are important for limiting meiotic germ cell losses.

### Estrogens deprivation impacts follicle formation

Another key feature of the phenotype of the *ARO^-/-^* females concerns the abnormal genital tract and ovarian follicle differentiation. At puberty, the genital tract of the XX *ARO^-/-^* rabbit was profoundly altered, with massive atrophy, as already reported in *Cyp19a1^-/-^* mice (Fisher *et al.* 1998; Britt *et al.* 2000), and in agreement with the fact already demonstrated that the development of the female genital tract is highly dependent on estrogens. The follicle differentiation that starts 12-15 days after birth in rabbit at the arrest of meiosis prophase is a continuous process that involves the surrounding of oocytes with flat pre-granulosa cells and the breakdown of ovigerous nests thus giving rise to primordial follicles. Then, the differentiation of pre-granulosa cells from flat-to cuboidal-form cells signs the differentiation of primary follicles. Of course, the size of the population of primordial follicles is linked to the number of oocytes, the latter being the consequence of all the events which occurred during fetal and neonatal life as discussed previously. In the *ARO^-/-^* rabbit ovary, this is of particular importance since the pool of oocytes and that of *FOXL2* expressing pre-granulosa cells are extremely reduced. More, it has been reported that the nests breakdown is an event specifically sensitive to apoptotic losses (Myers *et al.* 2014). This could explain why the pool of primordial follicles was very low in *ARO^-/-^* rabbit ovary few days after birth (Figure 11) and almost null at puberty (Figure 2). The poor expression of the *FOXL2* marker gene by the granulosa cells 16 days after birth signs the abnormal differentiation of the primary follicles, the number of which was extremely low (Figure 2). Surprisingly, controversial results were reported in *Cyp19a1* KO mice, where the number of primary follicles was significantly higher than in wild type mice (Britt *et al.* 2000). To date, we have no explanation for such differences. Furthermore, we observed that despite the absence of estradiol, large antral follicles with oocytes developed in the *ARO^-/-^* ovary, however these follicles were very few. This proves that estrogens are not the only actors in follicular growth and differentiation at puberty and onwards. Many previous studies have shown that these events depend on the interactions between several signaling pathways, among which are estrogens but also the anti-Müllerian hormone (AMH) and gonadotropins (Grynberg *et al.* 2012; Dewailly *et al.* 2016; Chou and Chen 2018). Additional studies are underway to further analyze the mechanisms of differentiation of these follicles in the absence of estradiol.

To conclude, the present work shows that in the rabbit species, the *CYP19A1* gene mutation and the absence of estrogens from early embryo life was at the source of a series of major disorders of the ovarian differentiation process. Two developmental stages were thus identified, where the autocrine or paracrine activity of ovarian estrogens is crucial for ovarian differentiation: i) a first stage during fetal life when estrogens act on cell proliferation and differentiation mainly on the coelomic epithelium, but also on germ cells to achieve the formation of ovigerous nests ii) a second stage after birth and germ cell meiosis, when estrogens act on ovarian follicle formation. Our hypothesis is that it is the same in other mammals when a long period with numerous mitoses intervenes between the determinism of the sex of the gonad and the induction of meiosis i.e. in several mammal species but not in mice. This highlights the importance of the fetal period and thus shows why the maternal environment in such species can have a major impact to induce irreversible effects. In addition, these data show that while estrogen therapy given after birth may have positive effects on the development of the genital tract and the metabolism, it cannot attenuate the effects of aromatase deficiency on early ovary differentiation.

## Materials and methods

### Animals

New Zealand rabbits (NZ1777, Hypharm, Rousssay, France) were bred at the SAAJ rabbit facility (Jouy-en-Josas, France). All experiments were performed with the approval of the French Ministry MENESR (accreditation number APAFIS#6775-2016091911407980 vI) following the recommendation given by the local committee for ethic in animal experimentation (COMETHEA, Jouy-en-Josas). All researchers working directly with the animals possessed an animal experimentation license delivered by the French veterinary services. Hormonal treatments for superovulation and surgery for embryo transfer were performed as previously described (Peyny *et al.* 2020).

### Design of TALEN sequences and plasmid constructions

Exon II of rabbit AROMATASE gene was targeted to create InDel mutations near the site of initiation of translation (ATG site). The target sequences (left arm: 5’-TGCTTCATCTGAAGCCA-3’ (sense); right arm: 5’-TGGGTTCAGTATTTCCA-3’ (antisense); 16 bases spaced) were chosen with the ZiFiT Targeter sofware (http://zifit.partners.org). No homology was identified at any other location in the rabbit genome that could represent a potential off-target site (supplementary table 1). The TALEN were constructed as described (Sander *et al.* 2011), see in supplementary methods). Each TALEN RNA was diluted (100 ng/μl) in injection buffer (Millipore, France) and stored at −80°C until used.

### Generation of mutant rabbits

Embryos produced from superovulated females were injected at single-cell stage with an equimolar mixture of the left and right arm TALEN mRNAs (50ng/μl each). Injected embryos were implanted 3-4 hours after injection into the oviducts of anesthetized recipient rabbits via laparotomy. For all details concerning handling of females and embryos, see in (Peyny *et al.* 2020).

Offspring were screened for the presence of InDel mutations using genomic DNA extracted from ear clips (Jolivet *et al.* 2014). First detection of founders and characterization of mutation were performed using one set of primers located far upstream and downstream the targeted ATG (F0/R0, Figure 1 and supplementary table 2 for all primer sequences). The amplified fragments were sequenced (Eurofins Genomics, Courtaboeuf, France) and the extent of the mutation was deduced by comparing with the sequence of a wild type rabbit.

For further routine screening of mutants, a quantitative PCR (Fast SYBR Green Master Mix, Applied Biosystems, ThermoFisher, France) was performed using two sets of primers, the *CYP19A1* specific one (*CYP19A1* gene specific primers) flanking the position of the characterized InDel mutations (F1/R1, supplementary table 2) and another set to amplify a 2 copies reference gene (the rabbit *beta GLOBIN* gene, ENSOCUG00000000568). By using the Δ(ΔCt) method with a wild type rabbit DNA as reference DNA, we deduced the copy number of the *CYP19A1* allele in each DNA sample (no amplification = the two alleles are mutant (*ARO^-/-^* genotype); one copy = one mutant allele (*ARO^-/+^* genotype); two copies = wild type (WT or *ARO^+/+^* genotype).

The presence/absence of the Y chromosome was deduced from the amplification of the *SRY* gene through PCR analyses with the rabbit *beta GLOBIN* gene as positive control of the PCR and *SRY* specific primers (supplementary table 2). All rabbits were tested to determine the genomic sex in parallel to the external and histological observations. In the present paper, mentions of the XY or XX genotype always refer to the PCR determination.

### Histological and immunohistological analyses

Immediately after sampling, gonads were immersed in Bouin’s fixative or paraformaldehyde (4% PAF in PBS 1x), fixed for 24 to 72 hours then paraffin embedded. Microtome sections of 5μm thickness were processed. Hematoxylin-eosin-saffron (HES) colorations were performed by the @Bridge platform (INRAE, Jouy-en-Josas) using an automatic Varistain Slide Stainer (Thermo Fisher Scientific). Sirius Red-Fast Green colorations were performed manually.

In situ hybridization (ISH) was performed using the RNAscope ISH methodology (ACD, Bio-Techne SAS, Rennes, France). Briefly, probes at around 1000nt long were designed and produced by the manufacturer in order to match to the full-length cDNA of interest, taking care to reduce any cross-hybridization with non-specific targets. The list of all synthesized probes is given in supplementary table 3. Hybridization was performed on 5μm sections from PAF fixed tissue using labelling kits (RNAscope 2.5HD assay-brown or -red, or RNAscope 2.5HD duplex chromogenic assay blue (conjugated to horse radish peroxidase) and red (conjugated to alkaline phosphatase)) as recommended by the manufacturer. The red labelling (Fast Red) was observed as visible or fluorescent signals. Hybridization was considered as positive when at least one dot was observed in one cell.

Immunohistological analyses were performed on 5μm sections from PAF fixed tissue. The supplementary table 4 recapitulates the antibodies used.

All colored sections (visible, or fluorescent ones) were scanned using a 3DHISTECH panoramic scanner at the @Bridge platform (INRAE, Jouy-en-Josas).

### RNA extraction and RT-qPCR analyses

Gonads from rabbit fetuses were collected and immediately frozen at −80°C. Total RNA from each gonad was extracted using the RNeasy^®^ MicroKit (Qiagen, France). Quantitative PCR was performed on reverse transcribed RNAs (High Capacity Reverse cDNA Transcription kit with the included set of random primers, Applied Biosystems, ThermoFisher, France). Several PCR sets of primers and analyses were those published previously (Daniel-Carlier *et al.* 2013). Other sets of primers are given in the supplementary table 2. Data from significant and specific amplifications only were reported, thus excluding all amplifications with too high or variable Ct values, and non-specific or multiple amplifications.

### Measurement of estradiol, testosterone and anti Mülllerian hormone levels in serum samples and fetal gonads

Estradiol and testosterone were assayed by GC/MS according to the protocol described by Giton et al. (Giton *et al.* 2015) with modifications (Devillers *et al.* 2019). Female rabbit serum from *ARO^-/-^* genotype was used as the matrix for calibrators and quality control (QC) standards after twice charcoal dextran treatments. Sample extraction and purification, derivatization and determination of estradiol and testosterone levels in serum samples and fetal gonads are described in details as supplementary information (see also supplementary Table 5) or can be provided upon request. Anti Müllerian hormone levels were determined in 50 μl aliquots of serum samples by using an ELISA kit (AMH GenII ELISA, with AMH Gen II calibrators and controls, Beckman Coulter, Villepinte, France) as previously described (Bourdon *et al.* 2018).

### Statistics

The statistical analyses were performed using the GraphPad Prism 7 Software (GraphPad Software Inc., La Jolla, CA, USA). Because of the small number of samples in groups, comparisons between values were made by the Mann-Whitney test for non-parametric values. A probability lower than 0.05 was required for significance.

## Acknowledgements

The authors would like to thank the staff of the facility (SAAJ, INRAE, Jouy-en-Josas) for the care of the rabbits and Julie Rivière and Marthe Vilotte (UMR GABI, INRAE, Jouy-en-Josas) for their assistance on the histological plateau (@Bridge platform) and for the access to the virtual slide scanner (MIMA2 platform). Special thanks go to Danielle Monniaux (UMR PRC, INRAE, Nouzilly) for the time she spent in enriching discussions and highly valuable comments. This study was supported by ANR grants (GENIDOV:ANR-09-GENM-009; ARGONADS: ANR-13-BSV2-0017; ARDIGERM: ANR-2020-CE14). The authors declare that they have no competing interests.

