## supplementary data and methods for "Fetal estrogens are not involved in sex determination but critical for early ovarian differentiation"

### Supplementary information

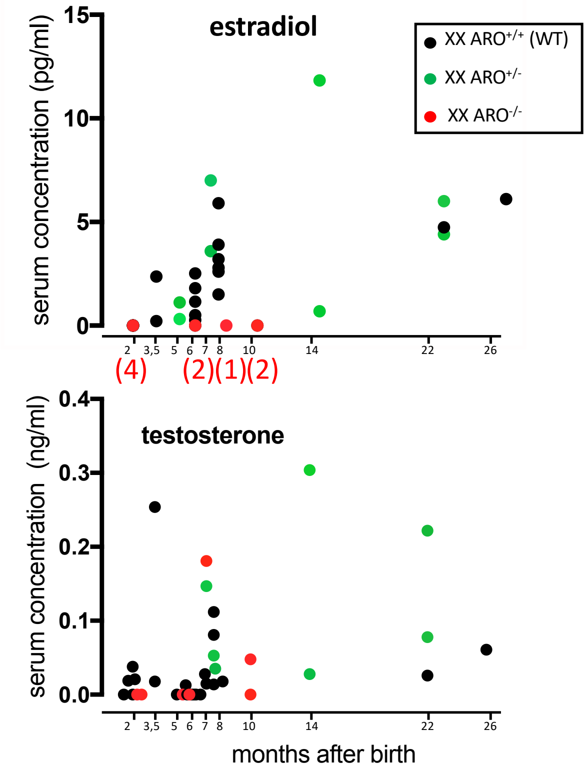


**Supplementary Figure 1**: **estradiol and testosterone serum concentrations in female rabbits collected from 2 to 26 months after birth.**

**Supplementary Figure 1**: **estradiol and testosterone serum concentrations in female rabbits collected from 2 to 26 months after birth.** Steroid concentrations were measured by GC/MS in serum of wild type XX rabbits (black points), *ARO*$^{+/-}$ rabbits (green points) and *ARO*$^{-/-}$ rabbits (red points). The number of analyzed sera samples from *ARO*$^{-/-}$ females is given in red between brackets. The same sera samples were used for assay of testosterone and estradiol concentrations.

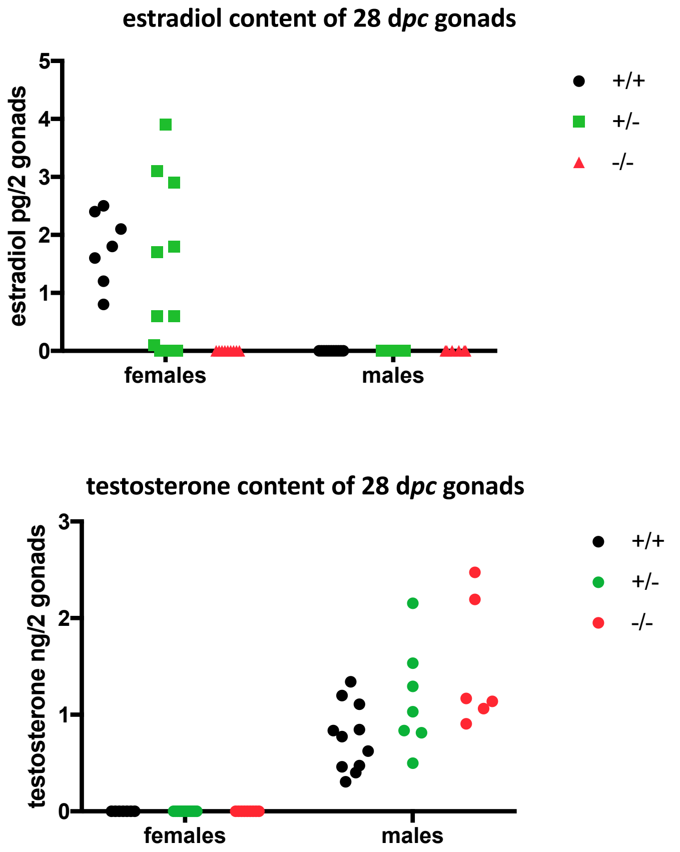


**Supplementary Figure 2: estradiol and testosterone content of 28 d*pc* gonads.**

**Supplementary Figure 2: estradiol and testosterone content of 28 d*pc* gonads.** Gonads of fetuses were collected 28 days after mating. Pairs of gonads were immediately frozen at -80°C. Steroid amounts were assayed by GC/MS. Each value is the amount of steroid in the pool of both gonads of one animal.

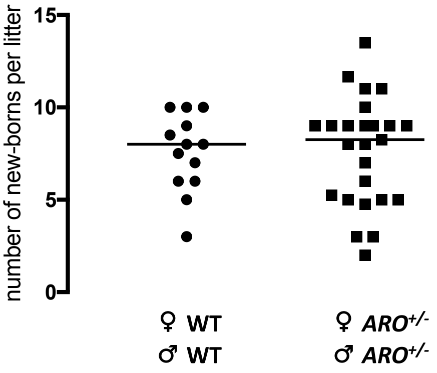


**Supplementary Figure 3**: number of neonates per litter after WT or *ARO*$^{+/-}$ mating

**Supplementary Figure 3: number of neonates per litter in WT or *ARO*$^{+/-}$ mating.** Mating were performed between *ARO*$^{+/-}$ males and *ARO*$^{+/-}$ females of the same mutant line, or between wild type males and females from the New-Zealand strain from the facility. Each point represents the mean size of litters from each female, each female having being mated by different males. When one female was mated several times by the same male, the graph reports the mean of the different litter sizes. The horizontal bars represent the medians.

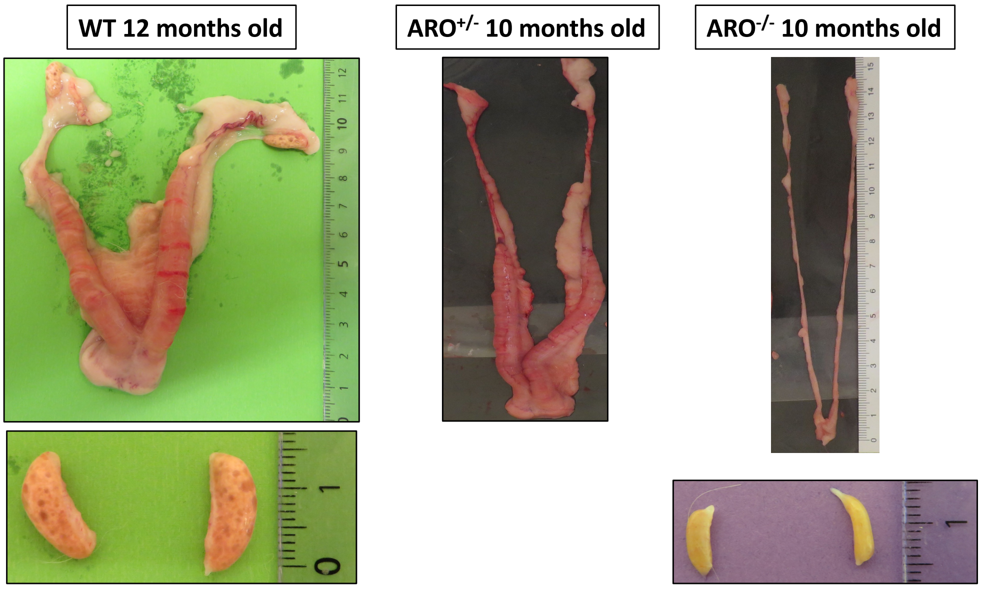


**Supplementary Figure 4: genital tracts of adult females**

**Supplementary Figure 4:** genital tracts of adult females from wild type (WT), heterozygous *ARO*$^{+/-}$ and homozygous *ARO*$^{-/-}$ animals (Δ339 strain). The pictures below show the ovaries from the same females after dissection.

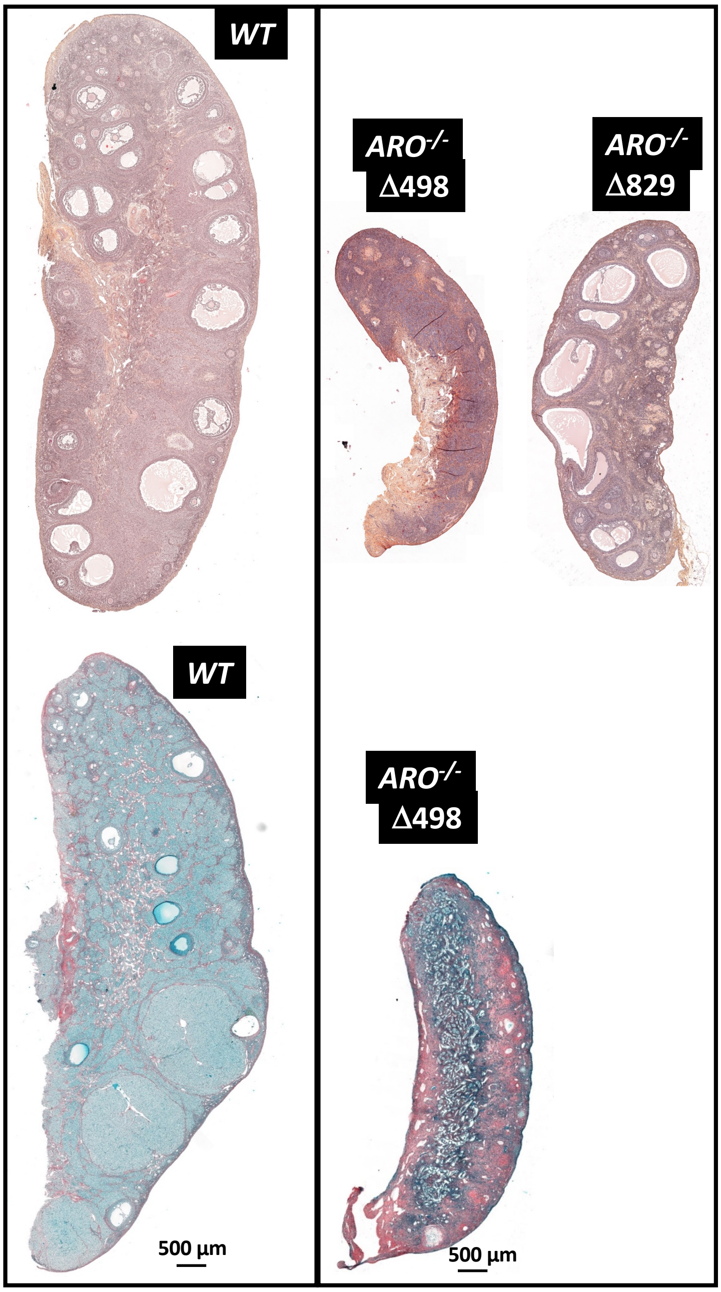


**Supplementary Figure 5: structure of ovaries of wild type (WT) and *ARO$^{-/-}$* rabbits from lines Δ498 and Δ829**

**Supplementary Figure 5: structure of ovaries of wild type (WT) and *ARO$^{-/-}$* rabbits from lines Δ498 and Δ829.**

The figure shows representative longitudinal sections of ovaries of 5-6 months old XX rabbits. Homozygous *ARO$^{-/-}$* XX mutants are issued from Δ498 and Δ829 lines. Organs were fixed with PAF then HES or Sirius red/fast green colored. With the Sirius-red/fast-green coloration (lower panel), collagen fibers were red colored. Cytoplasm was light-green colored and nuclei were dark-green. An intense red/collagen coloration was detected throughout the *ARO$^{-/-}$* ovary, with a strong staining in the cortical part. By contrast in the wild type ovary, collagen fibers were detected mostly in the medulla part. The Δ829 *ARO$^{-/-}$* ovary was filled with numerous large antrum follicles. However, the follicular reserve was almost null as in other *ARO$^{-/-}$* strains.

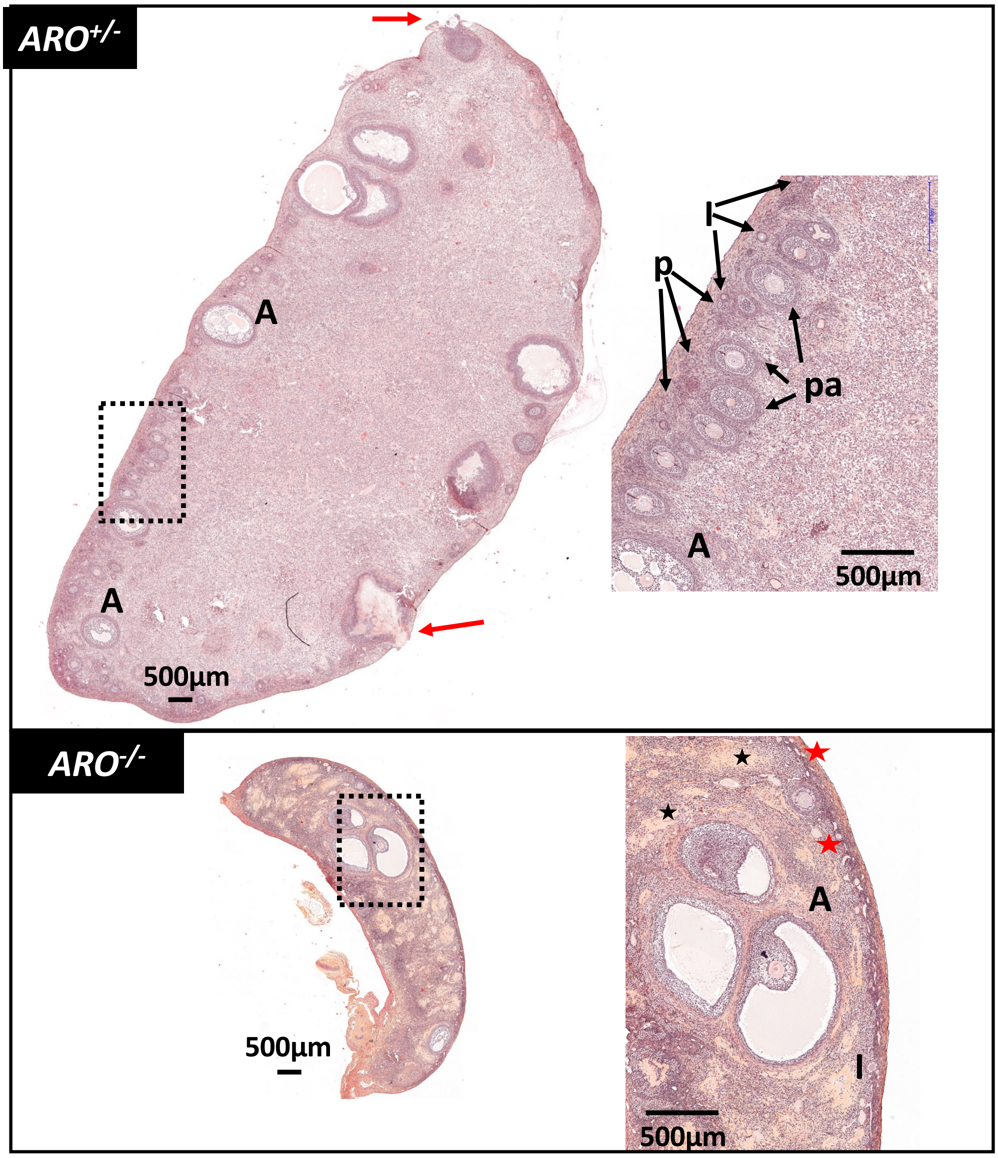


**Supplementary Figure 6. Ovaries of 10 months old rabbits after hormonal treatment to induce superovulation**.

**Supplementary Figure 6. Ovaries of 10 months old rabbits after hormonal treatment to induce superovulation**. Rabbits were hormonally treated for induction of ovulation, mated and sacrificed the following morning. Ovaries were collected and fixed (PAF) for histological HES colored observation. Follicular rupture (red arrow) occurred in *ARO*$^{+/-}$ ovaries, but not in *ARO*$^{-/-}$ ones. Antral (A), pre-antral (pa), primary (I) and primordial (p) follicles are pointed in enlargements. Black stars: accumulation of fibrous tissue.

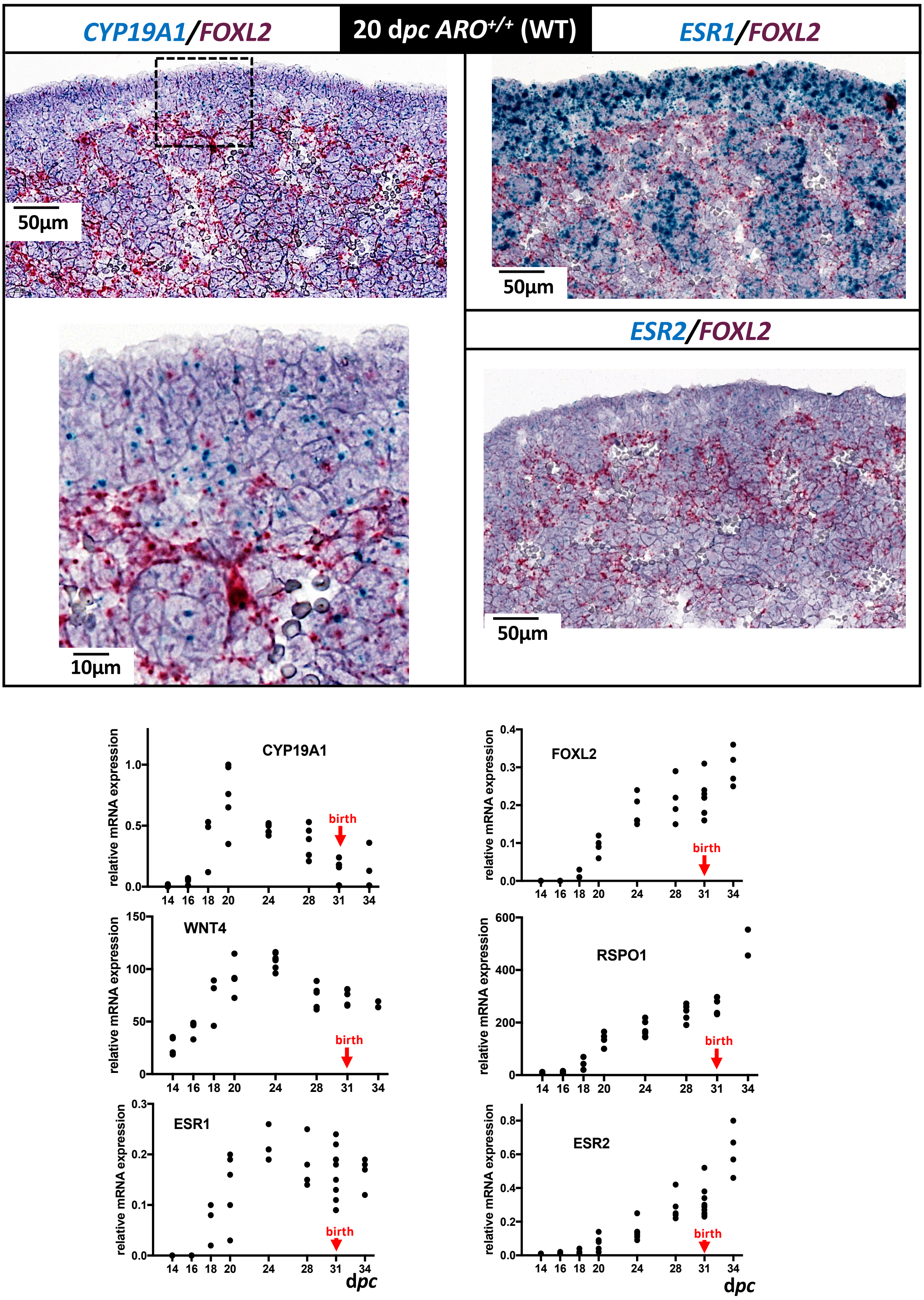


**Supplementary Figure 7:** ***CYP19A1, FOXL2, RSPO1, WNT4 and estrogen receptor*** **genes in wild type 20 d*pc* fetal ovaries**

**Supplementary Figure 7:** ***CYP19A1, FOXL2, estrogen receptor*** **genes in wild type 20 d*pc* fetal ovaries.**

The cellular localization of gene expression was determined using dual ISH with blue (*CYP19A1, ESR1, ESR2*) or red (*FOXL2*) labelling. Each picture represents the epithelium surface of the ovary and the tissue underneath. The black arrows point some cells with both blue and red signals in the enlarged picture.

The diagrams represent the relative mRNA expression levels. Each point represents one RNA sample extracted from both gonads of one animal at the indicated developmental stage. The same RNA samples were analyzed in the 4 diagrams. The vertical red arrow points the birth that occurs 31 days after conception in the rabbit species. Data concerning *CYP19A1* and *FOXL2* mRNA expression levels are those already published in Daniel-Carlier et al, 2013.

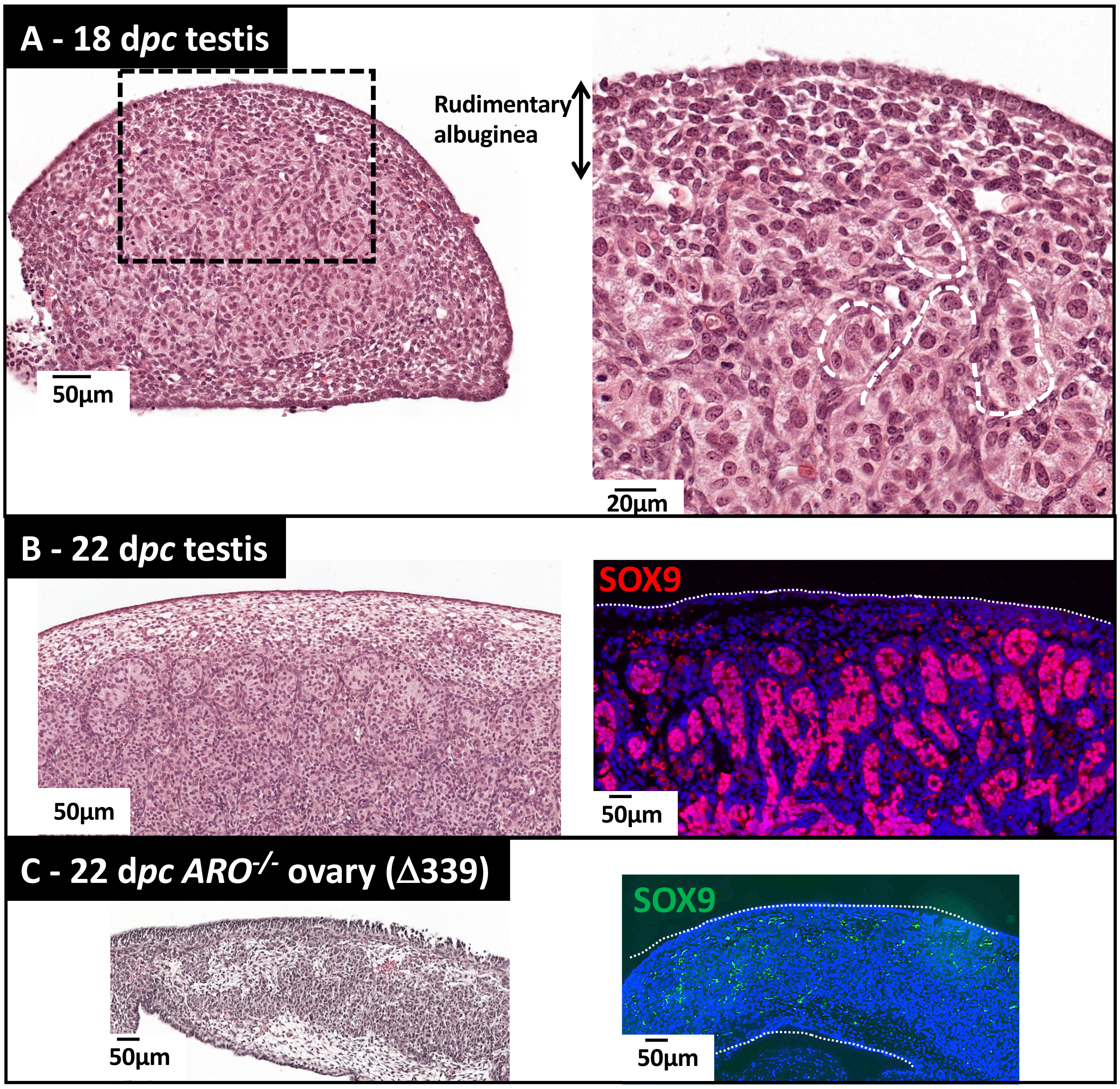


**Supplementary Figure 8: structure of 18 and 22 d*pc* old fetus testes**

**Supplementary Figure 8: structure of 18 and 22 d*pc* old fetus testes.** Gonads of rabbit fetuses from normal XY genotype were collected at 18 (**A**) and 22 d*pc* (**B, C**) then fixed in Bouin's. The picture in (**A**) shows a HES colored slice with an enlargement. The 18 d*pc* gonad developed as a testis, with differentiating seminiferous tubes (white dotted lines) wrapped in a rudimentary albuginea. The lower pictures (**B**, **C**) show HE staining (left panel) and fluorescence after labelling using an antibody directed against the SOX9 transcription factor (right panel). A wild type testis (**B**) and an *ARO*$^{-/-}$ ovary (**C**) were collected at 22 d*pc* then fixed in 4% PAF before immunohistological analysis. The SOX9 antibody (red fluorescence) labelled the nucleus of somatic Sertoli cells of the testis (**B**). No specific SOX9 labelling was detected in the *ARO*$^{-/-}$ ovary; the green fluorescent points correspond to non-specific labelling of red blood cells (**C**).

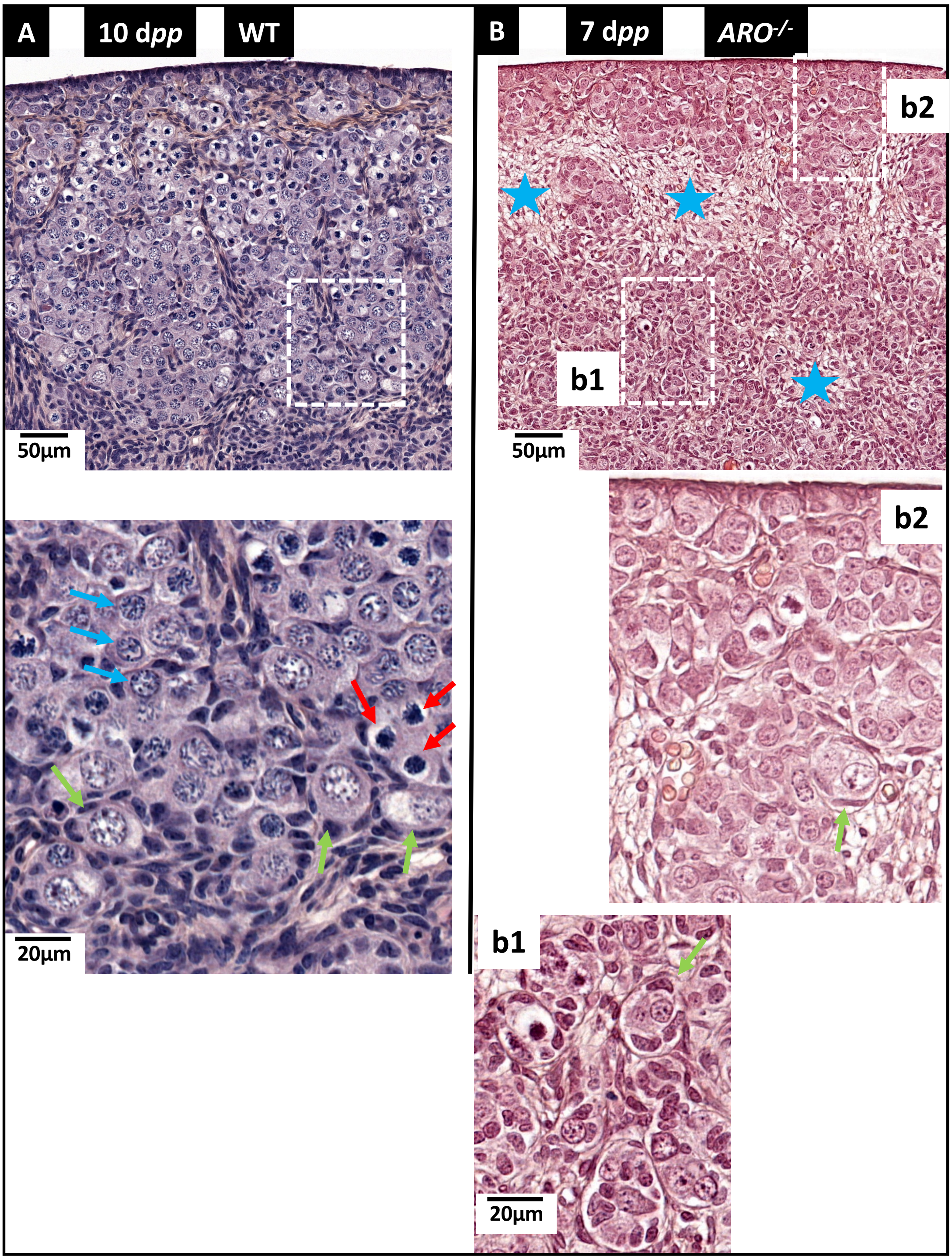


### Supplementary Figure 9: WT and *ARO*$^{-/-}$ ovaries seven days after birth

**Supplementary Figure 9: a few numbers of oocytes undergo the diplotene stage in *ARO*$^{-/-}$ ovaries seven days after birth**. Ovaries were collected 10 days (WT) or 7 days (*ARO*$^{-/-}$) after birth, treated by Bouin's fixative then paraffin embedded. Sections were HES colored. Blue stars point the connective tissue in *ARO*$^{-/-}$ ovaries. In wild type ovaries, red, blue and green arrows point the zygotene, pachytene and diplotene stages respectively. Somatic flat cells were found around some oocytes in both wild type (diplotene stage) and *ARO*$^{-/-}$ (green arrows, meiotic stage difficult to determine) ovaries.

| sequence of the targeted region TGCTTCATCTGAAGCCAAAGAGCACACGATGGTTTTGGAAATACTGAACCCA (52nt) | | | | |
|---|---|---|---|---|
| **Genomic location in the rabbit genome** | **overlapping gene** | **length of homology** | **% of homology** | **score** |
| 17:21282661-21282712 | CYP19A1 | 52 | 100 | 6,00E-21 |
| 1:41363894-41363914 | | 21 | 100 | 0.019 |
| 10:11913651-11913671 | | 21 | 100 | 0.019 |
| 19:31257617-31257637 | | 21 | 100 | 0.019 |
| 11:46433289-46433311 | | 23 | 95,65 | 0.29 |
| 14:61084103-61084121 | | 19 | 100 | 0.29 |
| 15:106972249-106972274 | | 26 | 92,31 | 1.2 |
| 16:27461959-27461980 | RYR2 | 22 | 95,45 | 1.2 |
| 2:70265022-70265039 | TUSC3 | 18 | 100 | 1.2 |
| 1:149838996-149839012 | TUB | 17 | 100 | 4.6 |
| 1:159223429-159223445 | | 17 | 100 | 4.6 |
| 1:171088368-171088384 | ELP4 | 17 | 100 | 4.6 |
| 1:75427837-75427853 | | 17 | 100 | 4.6 |
| 11:67725913-67725929 | | 17 | 100 | 4.6 |
| 12:55382223-55382239 | | 17 | 100 | 4.6 |
| 14:161219422-161219438 | | 17 | 100 | 4.6 |
| 14:78017712-78017728 | | 17 | 100 | 4.6 |
| 15:47467930-47467946 | SLC39A8 | 17 | 100 | 4.6 |
| 18:60262016-60262032 | VTI1A | 17 | 100 | 4.6 |
| 2:105241067-105241083 | | 17 | 100 | 4.6 |
| 2:172155957-172155973 | | 17 | 100 | 4.6 |
| 20:28226256-28226272 | | 17 | 100 | 4.6 |
| 20:31338072-31338088 | DNAL1 | 17 | 100 | 4.6 |
| 7:110853285-110853301 | | 17 | 100 | 4.6 |
| 7:117990691-117990707 | OSBPL6 | 17 | 100 | 4.6 |
| 7:168493075-168493091 | | 17 | 100 | 4.6 |
| 7:87316358-87316374 | | 17 | 100 | 4.6 |
| 7:91302331-91302347 | | 17 | 100 | 4.6 |
| 8:34242276-34242292 | M6PR | 17 | 100 | 4.6 |
| GL018700:7687914-7687930 | | 17 | 100 | 4.6 |
| GL018776:319476-319492 | | 17 | 100 | 4.6 |
| GL018909:412658-412674 | | 17 | 100 | 4.6 |
| X:34152874-34152890 | CLCN5 | 17 | 100 | 4.6 |
| X:52742450-52742466 | | 17 | 100 | 4.6 |

**Supplementary Table 1: Off-target sequences found by BLAST analysis as provided by the Ensembl genome browser.** The searched 52nt long sequence is given at the top of the table. The table reports the length of the homology sequence with the percentage and the final score. The best score (smallest number) was got, as expected, with the *CYP19A1* rabbit gene.

| gene name | sense primers | antisense primers | position of the amplified fragment in the rabbit genome or transcript |
|---|---|---|---|
| ***CYP19A1 (detection of mutants)*** | F0 5'-TGAATTCAACAGACAGCCTAATGG-3' | R0 5'-CCATTAGTGGTAGAATGGGAGG-3' | chromosome 17; -622, +864 relative to the translation initiation codon ATG of the *CYP19A1* gene (ENSOCUT00000012304) |
| ***CYP19A1 (routine genotyping)*** | F1 5'-TCTTCAATTTTTCTGCCCCTTTA-3' | R1 5'-TGCATTGGGTTCAGTATTTCCA-3' | chromosome 17; -71, +28 relative to the translation initiation codon ATG of the *CYP19A1* gene (ENSOCUT00000012304) |
| **rabbit $\beta$ *GLOBIN*** | 5'-GCCCCTCTGCTAACCATGTTC-3' | 5'-TTGCCAAAATGATGAGACAGCAC-3' | chromosome 1; +16,109; +16,200 relative to the translation initiation codon ATG of the $\beta$ *GLOBIN* gene (ENSOCUG00000000568) |
| ***SRY*** | 5'-GTTCGGAGCACTGTACAGCG-3' | 5'-GCGTTCATGGGTCGCTTGAC-3' | Y chromosome; +14, +158 relative to the translation initiation codon ATG of the *SRY* gene (Gene ID =100328958) |
| ***ESR1 = oestrogen receptor alpha*** | 5'-GCACCCAGGGAAGCTTCTAT-3' | 5'-AGCCAGCAACATGTCAAAGATTT-3' | chromosome 12; +1199;+1299 relative to the translation initiation codon ATG of the ESR1 cDNA (ENSOCUT00000004827.4) |
| ***ESR2 = oestrogen receptor beta*** | 5'-CTCACCAAGCTGGCTGACAA-3' | 5'-AGAGGCGCACTTGGTCCAA-3' | chomosome 20; +882;+982 relative to the translation initiation codon ATG of the ESR2 cDNA (ENSOCUT00000005082.4) |

**Supplementary Table 2 : Sequences of the primers used for genotyping**

The position of the amplicon is given relative to the translation initiation codon ATG of each gene. All primers match exactly with the sequence of the gene (wild type) and amplify a 100-140bp-long DNA fragment. Other primers are those used in the previously published paper (Daniel-Carlier *et al.* 2013).

| gene name | RNAScope probe catalogue number | transcript accession number (total length of cDNA in bp from ATG to STOP codon) | Position of probe versus ATG and STOP position |
|---|---|---|---|
| CYP19A1 (AROMATASE) | 440821 | NM_001170921.2 (1512bp) | 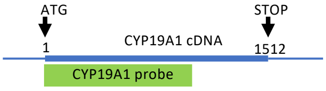  |
| DDX4 | 402341 | XM_002714 040.3 (2190bp) | 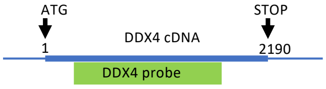  |
| ESR1 | 865461 | XM_002714947.3 (1797bp) | 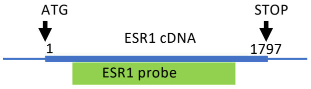  |
| ESR2 | 803171 | ENSOCUT00000 005082.3 (1572bp) | 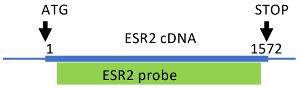  |
| FOXL2<br>FOXL2-C2 | 488221<br>488221-C2 | 3'UTR of a putative rabbit FOXL2 gene (985bp) | 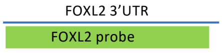  |
| OCT4<br>OCT4-C2 | 513271<br>513271-C2 | NM_001099 957.1 (1083bp) | 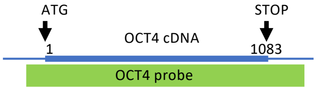  |
| RSPO2 | 803161 | ENSOCUT0000 0000779 (732bp) | 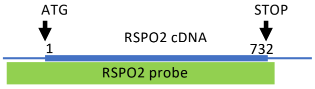  |
| SPO11 | 563501 | ENSOCUG0000 0000378 (1188bp) | 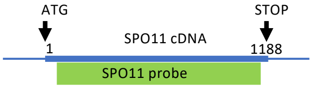  |
| STRA8 | 549411 | XM_008258 146.2 (1101bp) | 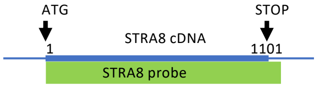  |

**Supplementary Table 3. In Situ Hybridization probes.** For each gene, the accession number of the transcript, the targeted region within the transcript and the catalogue number of the probe are given. The "C2" name refers to the probes conjugated to alkaline phosphatase; other probes are conjugated to horseradish peroxidase.

| protein | source | type | reference | working concentration |
|---|---|---|---|---|
| AMH | mouse | monoclonal (B-11 clone) | Santa Cruz sc-166752 | 1:50 |
| AROMATASE | mouse | monoclonal (H4 clone) (C-term) | ABD Serotec MCA2077S | 1:200 |
| γH2AX | mouse | monoclonal (D7T2V clone) | Cell Signaling | 1:200 |
| KI67 | rabbit | monoclonal (SP6 clone) | Thermo Fisher | 1:200 |
| OCT3/4 | goat | polyclonal | Santa Cruz Sc-8628 | 1:100 |
| SOX9 | rabbit | polyclonal | gift from Francis Poulat (IGH, Montpellier) | 1:200 |

**supplementary table 4** : antibodies and dilutions

**Supplementary material and methods:**

**Talen assembly**

The TALEN kit used for TALE assembly was a gift from the Keith Joung laboratory (Addgene kit # 1000000017). The left TALEN was constructed by assembling units of the kit in the following order: 9, 12, 20, 25, 27, 11, 20, 22, 30, 14, 16, 21, 29, 12, 17 and 21 (by groups of four units). The right TALEN was constructed by assembling units of the kit in the following order: 9, 14, 19, 25, 30, 12, 16, 24, 30, 11, 20, 25, 30, 12, 17 and 21. Each right and left insert was subcloned into the *Bsmb1* restriction site in JDS70 plasmid. Sequencing of the cloned DNA fragments was performed to check for the identity with what expected.

TALEN mRNAs were prepared from the TALEN-JDS70 plasmids using the ARCA T7 capRNA pol kit (Cellscript, TEBUbio France), polyadenylated with the polyA polymerase tailing kit (Epicentre Biotechnologies) then purified using the Qiagen RNEasy minikit (Qiagen France).

**Measurement of estradiol level in serum sample or gonads by Gas Chromatography / Mass Spectrometry (GC/MS)**

**Sample extraction and purification:** Briefly, samples (1 ml of serums, or the 2 gonads of one foetus, calibration standards, quality controls, and blank matrix) were collected in 8 ml borosilicate tubes. A spiking solution of deuterated steroid internal standard (IS) (50 µl containing 10 pg of E2-d4, except for blank matrix) (CDN Isotopes, Inc., Point-Claire, Canada), and 3 ml of 1-chlorobutane were added to each sample. Gonads were grinded using a small glass piston. After mixing (Vortex) and rapid centrifugation, the upper organic phase was collected and layered on conditioned Hypersep SI 500mg SPE minicolumn (Thermo Scientific, Rockwood, USA). The column and adsorbed material were then washed with ethyl acetate / hexane (6 ml; 1/9, v/v). The second fraction containing oestradiol was eluted using ethyl acetate / hexane (4 ml; 1/1, v/v), then evaporated at 60°C to dryness.

**Derivatization reaction and determination of rabbit plasma E2 level:** Estradiol was derivatized with pentafluorobenzoyl chloride (PFBC) (103772-1G, Sigma-Aldrich, Steinheim, Germany). Final extracts were reconstituted in isooctane, then transferred into conical vials for injection into the GC system (GC-2010 Plus, Shimadzu, Japan) using a 50% phenylmethylpolysiloxane VF-17MS capillary column (20m x 0.15mm, internal diameter, 0.15µm film thickness) (Agilent Technologies, Les Ulis, France). A TQ8050 (Shimadzu, Japan) triple quadrupole mass spectrometer equipped with a chemical ionization source and operating in Q3 single ion monitoring mode was used for detection. The reagent gas for the NCI detection was methane. The GC was performed in pulsed splitless mode with a 1 min pulsed splitless-time. The oven temperature was initially 150°C for 0.50 min, further increased to 305°C at 20°C/min and held at 305°C for 3.60 min, and then to 335°C at 30°C/min and held at 335°C for 1.7 min. The injection port and transfer line temperatures were respectively 290 and 280°C. The flow-rate of helium (carrier gas) was maintained constant at 0.96 ml/min. The mass spectrometer CI source temperature was 220°C. The linearity of steroid measurement was confirmed by plotting the ratio of the steroid peak response / internal standard (IS) peak response to the concentration of E2 for each calibration standard. Accuracy, target ions, corresponding deuterated internal control, range of detection, low limit of quantification (LLOQ), and intra & inter assay CVs of the quality control are given in the supplementary Table 5.

| **Accuracy** (%) | **Analyte** | **Target ion analyte / IS** (amu) | **Range** (pg/ml) $R^2$ | **Mean (17 runs)** ***Intra- & Inter assay CVs (%)*** | | | |
|---|---|---|---|---|---|---|---|
| | | | | **LLOQ Mean (pg/ml) Intra- & Inter assay CVs (%)** | **Low QC Mean (pg/ml) Intra- & Inter assay CVs (%)** | **Middle QC Mean (pg/ml) Intra- & Inter assay CVs (%)** | **High QC Mean (pg/ml) Intra- & Inter assay CVs (%)** |
| 94 - 107 | E2 | 660 / 664 | 0.2 - 56.0 | 0.22 | 2.87 | 6.07 | 12.88 |
| | | | 0.9994 | *17.3 - 19.6* | *2.9 - 4.2* | *2.6 - 3.7* | *2.3 - 3.3* |

**Supplementary Table 5: GC/MS plasma analytical control validation.**
*LLOQ : low limit of quantification*
*QC : quality control*
